# Marangoni-like tissue flows enhance symmetry breaking of embryonic organoids

**DOI:** 10.1101/2023.09.22.559003

**Authors:** Simon Gsell, Sham Tlili, Matthias Merkel, Pierre-François Lenne

## Abstract

During early development of multi-cellular animals, cells self-organize to set up the body axes, such as the primary head-to-tail axis, based on which the later body plan is defined. Several signaling pathways are known to control body axis formation. Here, we show, however, that also tissue mechanics plays an important role during this process. We focus on the emergence of a primary axis in initially spherical aggregates of mouse embryonic stem cells, which mirrors events in the early mouse embryo. These aggregates break rotational symmetry to establish an axial organization with domains of different expression profiles, e.g. of the transcription factor T/Bra and the adhesion molecule E-cadherin. Combining quantitative microscopy and physical modeling, we identify large-scale tissue flows with a recirculation component and demonstrate that they significantly contribute to symmetry breaking. We show that the recirculating flows are explained by a difference in tissue surface tension across domains, akin to Marangoni flows, which we further confirm by aggregate fusion experiments. Our work highlights that body axis formation is not only driven by biochemical processes, but that it can also be amplified by tissue flows. We expect that this type of amplification may operate in many other organoid and *in-vivo* systems.

## I. INTRODUCTION

Deciphering the mechanisms underlying the formation of the mammalian body plan is an important challenge in biology. This process takes place in the early embryo and is characterized by the establishment of a multicellular spatial organization with reference to an emerging system of orthogonal axes: the head-to-tail axis, also called the anteroposterior (AP) axis, and an orthogonal dorsoventral axis [1]. There is evidence that a small number of conserved morphogens, including Wnt, BMP, Activin/Nodal, and FGF, control the emergence of the body axes [2]. These signals participate in tissue patterning and trigger differential behaviors, which ultimately partition an isotropic group of cells into different territories. Yet, the physical mechanisms underlying axis formation remain largely unknown.

Here we focus on the emergence of the head-to-tail body axis, which breaks the initial rotational symmetry. In vertebrates, the AP axis establishes first by locating the expression of an evolutionarily conserved transcription factor, T/Brachyury (T/Bra), to one end of the embryo. Yet, understanding the emergence of the AP axis, and the appearance of the T/Bra pole in particular, is a daunting task because of the experimental difficulty in accessing the embryo, in particular in mammals [3– 5]. To address this issue, an excellent experimental system is provided by mouse and human embryonic stem cells (ESCs). 3D spherical aggregates of mouse or human ESCs self-organize into gastruloids, which are embryonic organoids that undergo symmetry breaking and exhibit axial organization and gene expression patterns that mirror events in the embryo [6–10]. Depending on the differentiation protocol, i.e. depending on the culture conditions and the signaling molecules present in the medium, gastruloids have the capacity to develop advanced structures strikingly similar to organs such as (1) somites and neural-tube [11–14] or (2) gut and heart [11, 15, 16]. Past works on gastruloids have identified the temporal sequence of gene expression [7] and networks of genes that regulate the main steps of their early development, including during symmetry breaking [17].

Yet, there is growing evidence that tissue mechanics and flows are crucial for many stages of embryonic development [18–22]. Indeed, in avians it has recently been pointed out that tissue mechanics and flows could also contribute to embryonic axis formation [23–25], but their potential role for axis formation in mammals remains unclear.

At least for one differentiation protocol, we have recently shown that symmetry breaking in mouse gastruloids is accompanied by large-scale tissue flows and heterogeneities in cell-cell adhesion and cell differentiation [16]. In this protocol derived from the standard protocol in the field [26], we first seed a controlled number of pluripotent mouse ESCs into non-adherent micro-wells (Figure 1a, Methods). After a few hours, the cells form 3D aggregates. 48 hours after aggregate formation (hAAF), differentiation is stimulated by adding the Wnt-agonist Chiron, which is washed out 24 hours later. Note that in this manuscript, following the usual convention in the field, when indicating time points in hAAF, the time of aggregate formation refers to the time when the cells were seeded. At the end of Chiron exposure (72 hAAF), gastruloids are spherical and express T/Bra throughout. Between 72 and 96 hAAF, gastruloids start to elongate and display AP-graded T/Bra and E-cadherin (E-cad) expression patterns with higher expression posteriorly (Figure 1a-d). These heterogeneities are linked to an epithelial-to-mesenchymal transition (as evidenced by the differential expression of Snail, Figure 1d, Supplementary Figure 1), which has since been identified in other differentiation protocols [27]. The main tissue types produced after symmetry breaking in our differentiation protocol are endoderm and mesoderm ([16], Supplementary Figures 2-5).

**FIG. 1.**
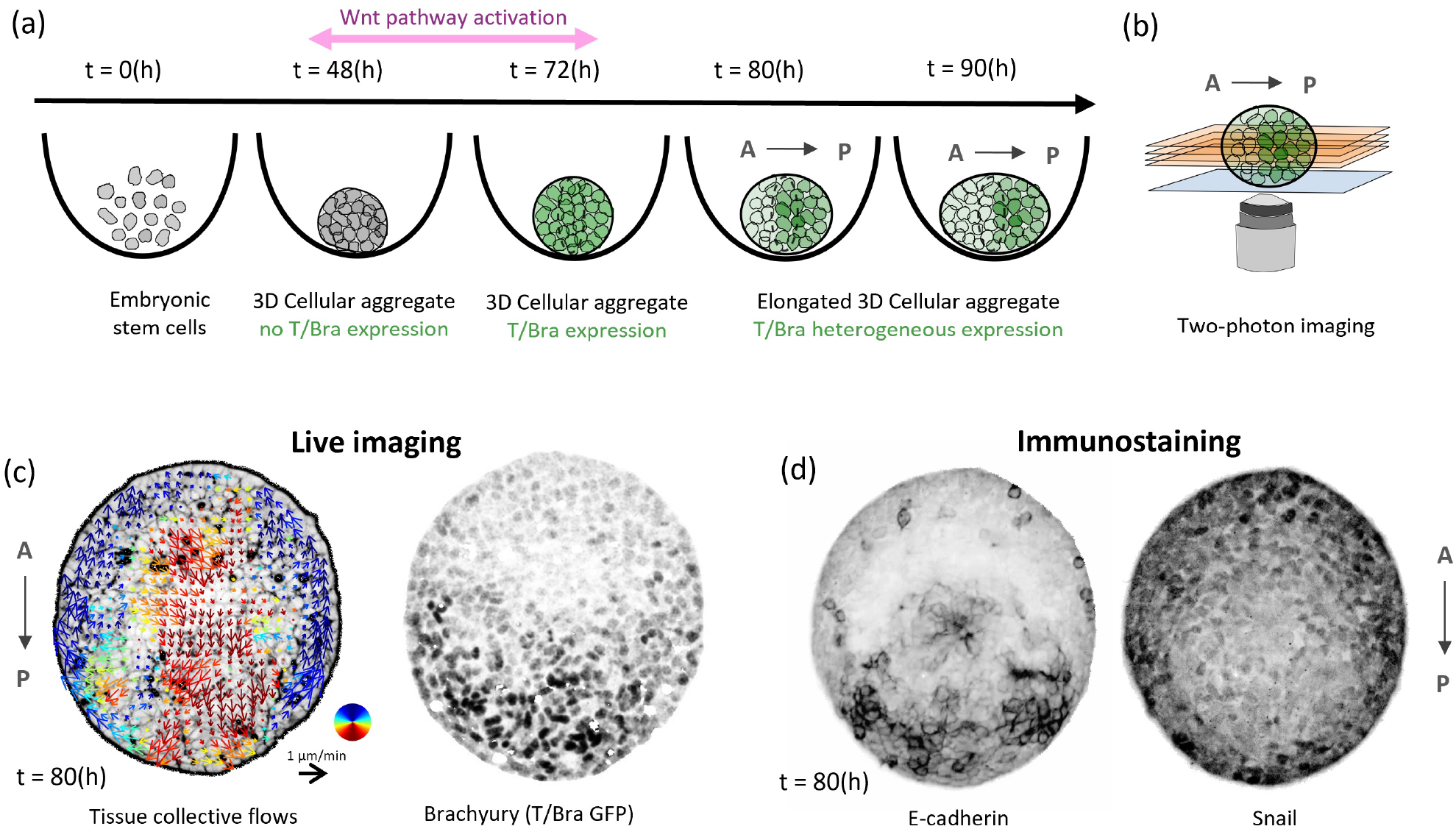
Experimental protocol to study gastruloid symmetry breaking. (a) Protocol to form gastruloids in low-adherence microwells. (b) Inverted two-photon microscopy is used to image sub-volumes of the gastruloids. (c) From live imaging data, we obtain tissue flows associated to gastruloid symmetry breaking (averaged over one hour) and the corresponding T/Brachyury pattern (gray levels are inverted for better visibility). (d) E-cadherin and Snail patterns obtained by immunochemistry, same gastruloid at a similar time as in panel c (gray levels are inverted for better visibility).

Here we quantitatively show that large-scale tissue flows contribute substantially to symmetry breaking in our mouse gastruloids. We show that the dominating component of the tissue flows is a recirculating flow whose direction correlates strongly with the T/Bra and E-cad polarization of the aggregate. We then present a mini-mal effective model for the emergence of these flows. In particular, we show that effective interfacial tensions between low and high T/Bra or E-cad domains, as well as differential surface tensions of these domains, are sufficient to explain the recirculating flows. Finally, we further validate our hypothesis of differential tensions by measuring interface angles in fusion experiments between gastruloids. Taken together, our work quantitatively shows that not only genetic and biochemical interactions, but also tension-driven large-scale tissue flows promote symmetry breaking in gastruloids.

## II. RESULTS

### Tissue flows substantially contribute to gastruloid symmetry breaking

In previous work, we provided experimental evidence for the existence of large-scale tissue flows, which differently advected cells with different levels of T/Bra or E-cad expression [16]. Motivated by these results, here we aim to precisely quantify the contribution of such tissue flows to the overall symmetry breaking. To this end, we need to simultaneously measure tissue flows and T/Bra or E-cad expression patterns.

We generate the gastruloids following our previously-published protocol (Figure 1a, Methods) [16], where we focus on the time between 72 and 96 hAAF, when gastruloids start to elongate and display polarized T/Bra and E-cad levels. Quantifying tissue flows and expression patterns in the gastruloids is challenging, as gastruloids are 3D cell aggregates with a diameter of a few hundred micrometers, which scatter light and are sensitive to prolonged illumination. To reduce light scattering while preventing phototoxic effects, we have used two-photon microscopy (see Methods) and imaged 60 µm-thick sub-volumes around the mid-plane of the gastruloids (Figure 1b, Methods). We ensured that the T/Bra and E-cad polarization direction, as well as the flow field directionality was approximately aligned with the imaging plane. To reduce drift during live-imaging, we transfer the gastruloids into arrays of 500 µm microwells at 72 hAAF (see Methods). We have used two cell lines: one with a T-Bra GFP live reporter, and one with an E-cad GFP live reporter (see Methods). To quantify tissue flows, we also added one of two different live cell markers, one for cell membranes and extra-cellular space (Sulforhodamin B, see Supplementary movie 1), and one for cell nuclei (SPY555-DNA; see Supplementary movies 2-5).

To quantify flow fields, we first applied a registration removing any global aggregate translation and rotation (see Methods), and then applied an optical flow method [28] to the resulting movie (see Methods, Supplementary Figure 6-7, Supplementary movie 6).

To quantify the degree of symmetry breaking of a gastruloid, i.e. its polarization, we use the dipole moment of its T/Bra or E-cad distribution (Figure 2a-c). It is represented at each time point *t* by the 2D vector ***P***, which we define as:

**FIG. 2.**
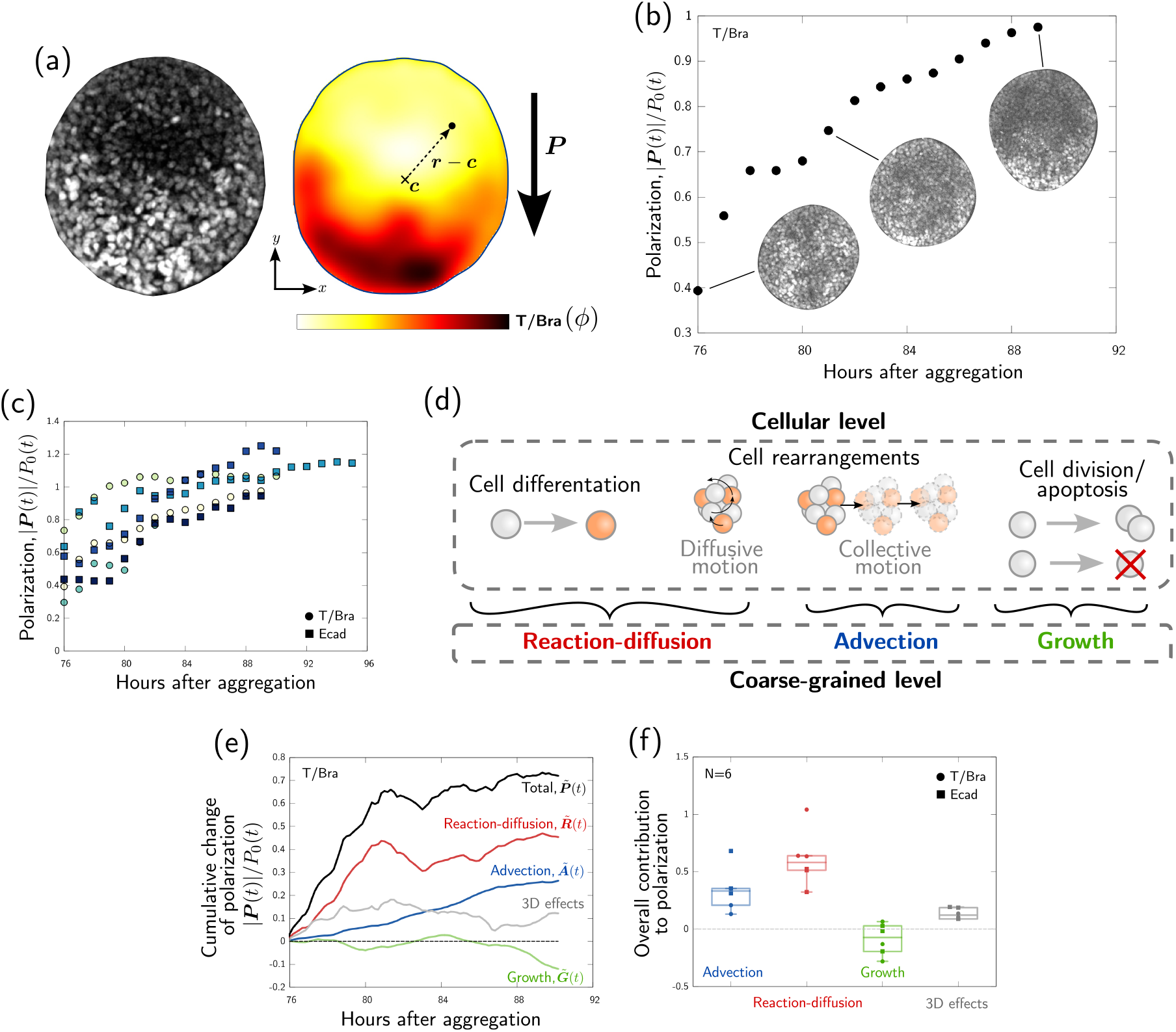
Quantitative analysis of the polarization process. (a) Example of T/Bra distribution and corresponding coarse-grained field *ϕ*. (b) Example of the time evolution of the T/Bra dipole moment and corresponding T/Bra distributions. (c) Time evolution of T/Bra and E-cad dipole moments for different samples. Each sample is indicated with a distinct color. (d) At the cellular level, the polarization process relies on the interplay between cell differentiation, cell rearrangements and cell division/apoptosis, resulting in three effective mechanisms at the coarse-grained level, namely reaction-diffusion, advection and growth. (e) Example of time evolution of the cumulative contributions to T/Bra polarization. For instance, 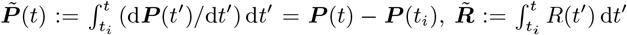 etc (see SI). (f) Relative total contribution of each process at the end of the polarization.

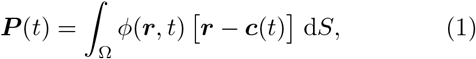

where Ω represents the 2D cross section of the aggregate, *ϕ* denotes the T/Bra or E-cad fluorescence intensity field, ***r*** is the position vector that we integrate over, and ***c*** is the barycenter of the aggregate. The direction of the vector ***P*** points towards the part of the gastruloid that has the highest fluorescence intensity in T/Bra or E-cad, and its magnitude quantifies the degree of symmetry breaking (i.e. polarization).

In our plots, we normalize the polarization vector ***P*** (*t*) at each time point by some reference value *P*_0_(*t*) that depends on the instantaneous spatial standard deviation of the T/Bra or E-cad expression level and the total cross-sectional area of the gastruloid (Figure 2b,c, see SI). This allows us to compensate for both time dependence and sample variability of overall expression level and size when comparing different gastruloids. Compared to a time-independent normalization, this resulted in a better collapse of the polarization dynamics across gastruloids (Figure 2c and Supplementary Figure 8b).

We found that after registration, the direction of the dipole moment does not change much over time (Supplementary Figure 8a). This indicates that very early processes already define an axis that is kept later on. However, we also see that the magnitude of ***P*** increases over time, reflecting a progressive build up of gastruloid polarization (Figure 2b,c).

We wondered how important different cellular processes are for the progressive increase in gastruloid polarization over time. There are different cell-scale processes that might contribute, which include (i) spatially heterogeneous cell differentiation, (ii) individual cell motion, (iii) collective cell motion (i.e. tissue flows), and differential (iv) cell division and/or (v) apoptosis (Figure 2d). To differentiate the contribution of all these processes to aggregate polarization, segmentation and tracking of the cell nuclei would be required, which is currently difficult due to light scattering and dense nuclear packing. Thus, we base our analysis here on coarse-grained optical flow and fluorescence data, which still allows us to differentiate three different contributions to gastruloid polarization: (1) the combined effect of cell differentiation, and individual cell motion (contributions i and ii), which we call for brevity “reaction-diffusion” (RD) contribution, (2) advection of T/Bra or E-cad with tissue flows (contribution iii), and (3) differential volume growth, arising for instance from cell division and apoptosis (contributions iv and v), which we call for brevity just “growth” contribution (Figure 2d).

Using the optical flow and fluorescence data, we can quantify the contributions 1-3 to the overall increase in polarization. To this end, we decompose the time derivative of ***P*** into four parts (Figure 2e, see SI):

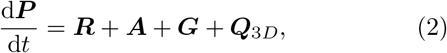

with

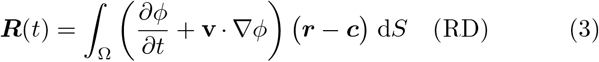

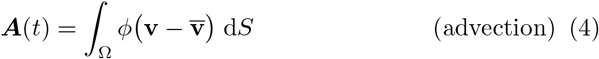

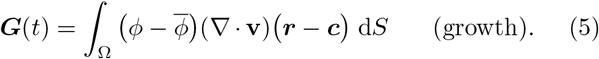

Here, **v** is the local tissue velocity (Figure 1c) and a line over a symbol,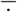, denotes averaging over the aggregate cross section. The first three terms on the right-hand-side of Eq. (2) respectively represent the contributions 1-3 (Figure 2e), and ***Q***_3*D*_ is a geometric contribution related to 3D motion of the aggregate (see SI). Besides this geometric contribution ***Q***_3*D*_, our mathematical decomposition also confirms that there is no other contribution to gastruloid polarization, as defined by Eq. (1), than contributions 1-3.

For all samples, we quantify each term in Eq. (2) over time and plot their respective cumulative contributions to the overall dipole moment. For one gastruloid, this is shown in Figure 2e, and for the other gastruloids in Supplementary Figures 8c-l. We compare the final overall contributions across all gastruloids in Figure 2f. We find that reaction-diffusion processes generally contribute most to the overall polarization. However, our analysis also reveals a significant contribution of advection, which is responsible for approximately 1/3 of the total polarization.

Taken together, our analysis shows that advection with large-scale tissue flows significantly contributes to gastruloid symmetry breaking. To better understand how, we next analyzed the structure of the flow pattern across different gastruloids.

### Tissue flows exhibit a dominant recirculating component

Qualitatively, tissue flows emerging during the polarization process exhibit substantial fluctuations, both spatio-temporally and across different gastruloids (see Supplementary Figures 7 and 9-12). In order to capture the dominant large-scale flow structures and compare them across gastruloids, we decomposed the tissue flow field into different modes (Figure 3 and SI). To this end, we define a disk that encompasses most of the aggregate projection (Figure 3a), and express the coarsegrained tissue velocity **v**(***r***) in polar form, i.e. with radial and angular velocity components v_*r*_(***r***) and v_*θ*_(***r***) (Figure 3b). We then decompose both spatial functions v_*r*_(***r***) and v_*θ*_(***r***) into different modes, again using polar coordinates. To capture the variation of both functions with the angular coordinate *θ*, we decompose them into angular Fourier modes (mode index *n*, Figure 3c). Similarly, we decompose the radius-dependent part of both functions into polynomial modes (mode index *p*, Figure 3c).

**FIG. 3.**
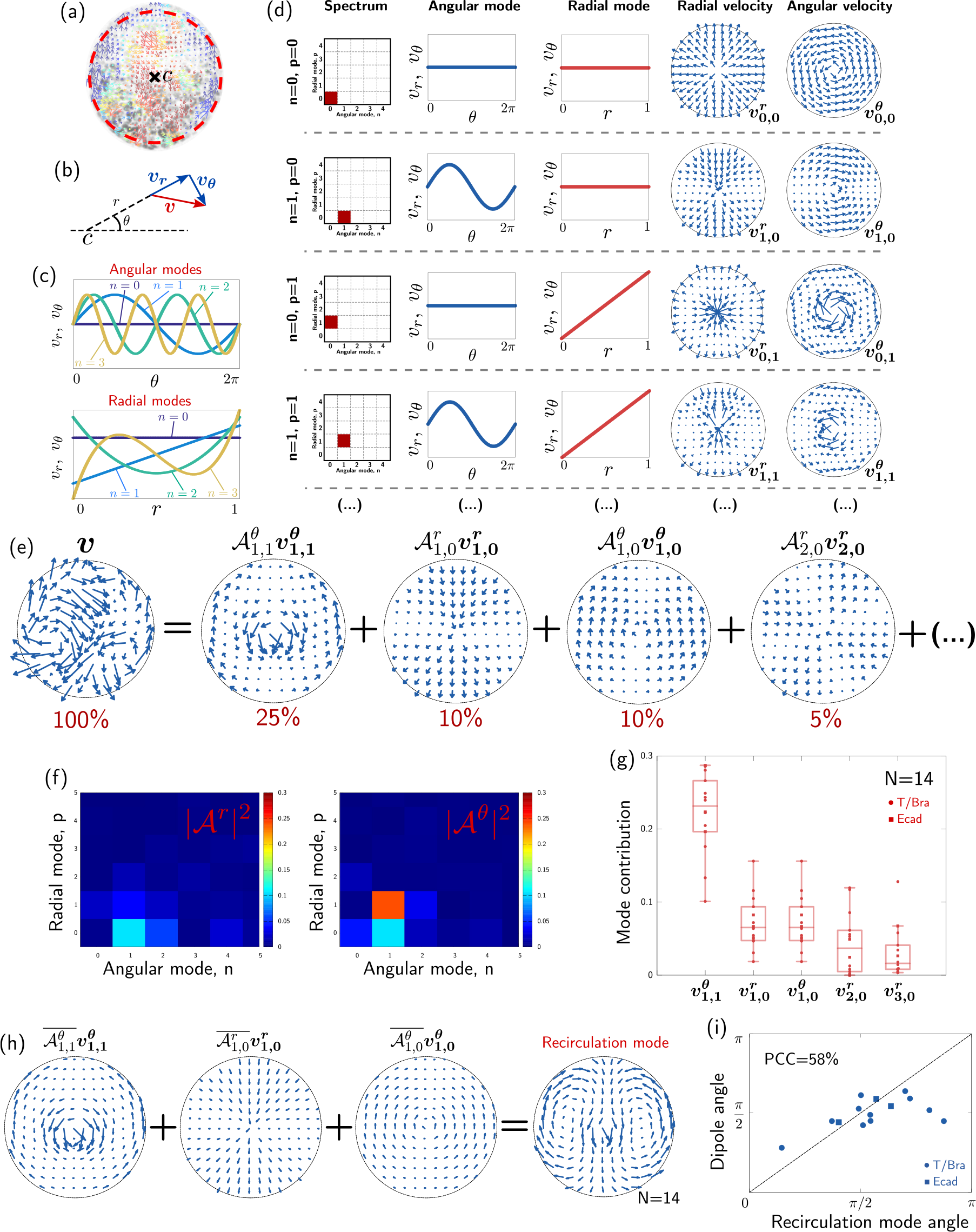
Analysis of tissue flow during polarization. (a) Coarse-grained tissue flows are analyzed in a circular disk, centered on the aggregate barycenter ***c***. (b) Tissue velocity **v** is decomposed as the sum of a radial component v_*r*_ and an angular component v_*θ*_. Similarly, positions within the aggregate can be expressed in polar coordinates (*r, θ*). (c) The velocity components v_*r*_(***r***) and v_*θ*_(***r***) are decomposed into a series of angular (index *n*) and radial (index *p*) modes. These modes are plotted for *n* ≤ 4 and *p* ≤ 4. (d) The first few modes are illustrated by their spectrum, the corresponding angular and radial functions, and the resulting radial and angular velocity fields. (e) Example of tissue velocity field expressed as a sum of the base modes in panel d, where the modes are weighted by the amplitudes 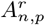 and 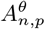. Below each mode we indicate the approximate contribution to the total variance of the flow. (f) Spectra displaying the contributions of each mode to the overall flow (compare panel d, leftmost column). Same sample as in (e). (g) Leading mode contributions over different samples. (h) We reconstruct the dominant recirculation mode over samples using median amplitudes of the three leading modes 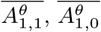 and 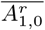. (i) Scatter plot showing the orientation angle of the recirculation mode versus the orientation angle of the dipole moment for each sample.PCC indicates the Pearson correlation coefficient.

As a result, both v_*r*_(***r***) and v_*θ*_(***r***) are decomposed into a sum of base modes 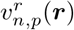 and 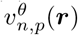, each weighted by a complex amplitude, denoted 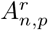 and 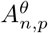:

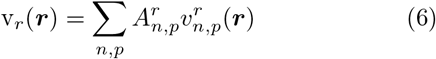

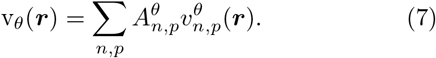

We display a few of the base modes 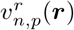 and 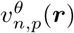 in Figure 3d. Generally, the modes with smaller indices (*n, p*) correspond to larger-scale features of the flow field. Each of the complex mode amplitudes 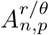 provides information about an angle and a magnitude. The angle corresponds to a rotation of the respective base mode around the barycenter, and the squared magnitude reflects the contribution of the respective base mode to the spatial variance of the flow field (Figure 3e,f; see SI). In other words, the modes with the largest amplitudes correspond to the dominant features of the flow field.

We apply our mode decomposition analysis on 9 gastruloids imaged between 78hAAF and 88hAAF. The analysis reveals that about 50% of the tissue flow field is provided by only 5 low-index modes, i.e. large-scale features, which are robustly observed across different gastruloids (Figure 3e-g). In particular, the three modes of highest amplitude, which are 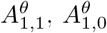 and 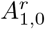, to-gether contribute approximately 40% to the total flow field. The superposition of these three modes corresponds to a large-scale recirculating pattern (Figure 3h). The next most relevant mode identified in Figure 3g, 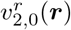 with amplitude 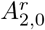 (Figure 3e right), contributes to gastruloid elongation. It corresponds to a flow field that extends along one axis and contracts along the perpendicular axis, consistent with gastruloid elongation between 72 and 96hAAF.

We find that the direction of the large-scale recirculating pattern strongly correlates with the T/Bra polarization, where the direction of the flow in the aggregate center points towards high T/Bra concentrations (Figure 3i). This correlation between the directions of T/Bra polarization and recirculating flow could be created by the advection of T/Bra with the tissue flows. However, could it also be possible that, conversely, aggregate polarization drives the recirculating flows?

### The recirculating flows could be driven by patterning-induced tension

To understand how the aggregate polarization – in terms of gene expression patterns – could affect tissue mechanics and flows, we first noted that from a coarse-grained perspective, the expression patterns of many genes are qualitatively similar in showing a distinct AP gradient (Figure 1c,d and Supplementary Figures 1-5). We thus studied a minimal model that represents these AP patterning genes by a single effective expression pattern *ϕ*(***r***), which ranges from −1 (anterior-most) to +1 (posterior-most). The field *ϕ*(***r***) reflects the combined effect of the AP patterning genes on tissue flows. In particular, the AP patterning genes include differentially expressed cadherins such as E-cad [16], which are known to lead to tension-driven phase separation [29–33].

To understand whether the observed gastruloid patterning can drive the recirculating flow, we study a simple continuum model that describes the gastruloid as an incompressible viscous fluid whose flows are driven by gradients of the field *ϕ*(***r***). Formally, this is described by Stokes’ equations:

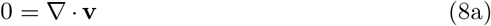

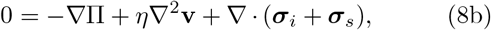

where Π is the hydrodynamic pressure ensuring incompressibility, *η* is the effective tissue viscosity, and ***σ***_*i*_ and ***σ***_*s*_ are symmetric, traceless tensors that denote additional mechanical stresses driving tissue flows. The stress ***σ***_*i*_ is internally created by the effective expression pattern *ϕ*(***r***), and ***σ***_*s*_ represents the tension at the aggregate surface.

There are different ways in which the internal stress ***σ***_*i*_ could depend on *ϕ*(***r***). However, as pointed out previously [34], the leading-order effect that drives incompressible flow is given by

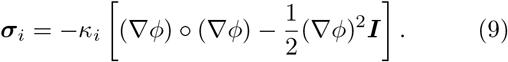

Here, the coefficient *κ*_*i*_ controls the magnitude of the internal mechanical stress ***σ***_*i*_, the symbol ° denotes the dyadic product, and ***I*** denotes the identity tensor. The internal stress tensor ***σ***_*i*_ is locally aligned with the gradient of *ϕ*(***r***), and its magnitude increases monotonically with the magnitude of the gradient (Figure 4a). At an interface between regions of low and high *ϕ*(***r***), the tensor ***σ***_*i*_ determines an effective interface tension *γ*_*i*_, which is obtained by integrating ***σ***_*i*_ across the interface width (Figure 4a,b).

**FIG. 4.**
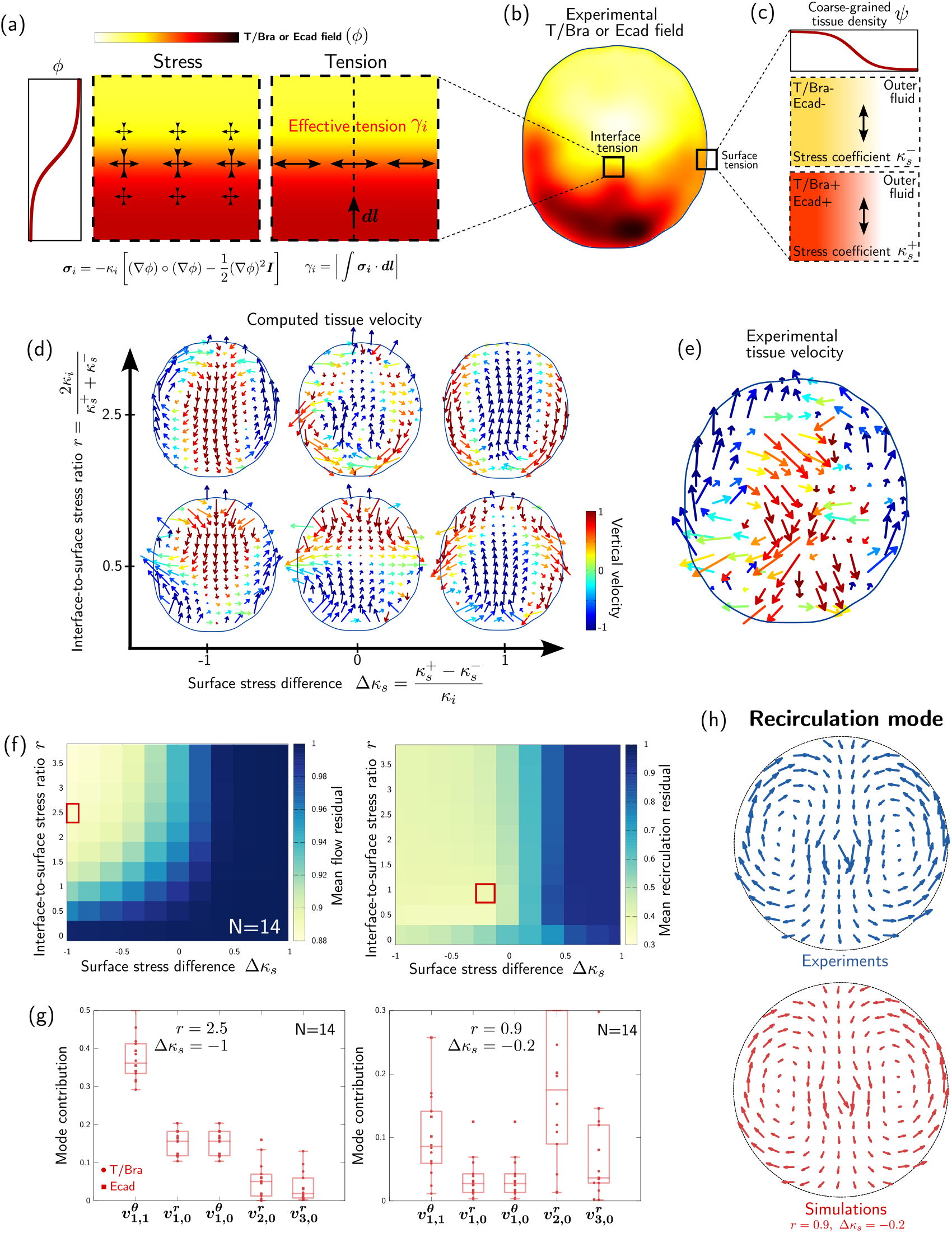
Computational modelling of the recirculating tissue flows. (a) Gradients of a scalar field *ϕ* control stresses, which results in an effective tension *γ*_*i*_ at the interface between domains of high and low *ϕ*. (b) We construct coarse-grained scalar fields from the experimentally observed T/Bra or Ecad fields and we compute the resulting active stresses within the aggregate. (c) We describe the aggregate surface as a smooth interface using a second scalar field *ψ*. The effective tension at the aggregate surface is controlled by a *ϕ*-dependent stress coefficient, with two limit values 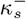 and 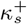 corresponding to surfaces of T/Bra (or E-cad-) and T/Bra+ (or E-cad+) tissues, respectively. (d) We compute tissue flows driven by the T/Bra- or E-cad-dependent stresses over ranges of the interface (*κ*_*i*_) and surface (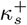 and 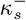) stress coefficients, varied through the non-dimensional parameters *r* and Δ*κ*_*s*_. The simulated flow fields shown here were computed from the T/Bra field plotted in panel b. (e) Experimental velocity field corresponding to panels b and d. (f) Residual between simulated and experimentally observed flow field, averaged over all gastruloids, as a function of the model parameters Δ*κ*_*s*_ and *r*. (left) Residual between the full flow fields, (right) residual between the modes contributing to the recirculation flow only. (g) Dominant modes contributing to the simulated flow field for two parameter sets, respectively minimizing the total flow residual (*r* = 2.5, Δ*κ*_*s*_ = −1) and the recirculating flow residual (*r* = 0.9, Δ*κ*_*s*_ = −0.2). (h) Reconstructed recirculation flow in experiments and simulations.

To describe aggregate surface tension ***σ***_*s*_, we introduce a second field *ψ*(***r***), which is equal to 1 inside the aggregate and 0 outside (Figure 4c). The stress ***σ***_*s*_ can be computed from gradients of *ψ*(***r***) analogously to Eq. (9), but with a magnitude *κ*_*s*_ that depends on local cell fate, i.e. on the *ϕ*(***r***) field. In the two limiting cases of *ϕ* =−1 (anterior regions) and *ϕ* = +1 (posterior regions), we denote the surface stress magnitude by 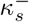 and 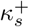, re-spectively (Figure 4c, details in the SI).

To test how far our simple effective model can account for the observed tissue flows, we solve Eqs. (8) and (9) using a lattice Boltzmann approach [35, 36] (see details in the SI), where we use for *ϕ*(***r***) the coarse-grained experimentally determined T/Bra or E-cad fields, as representative AP expression patterns. In Figure 4d, we plot for one of the T/Bra fields the resulting tissue flow field varying two dimensionless stress ratios, a relative surface stress difference, 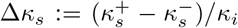 and a interface-to-surface stress ratio, 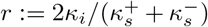. We find that the surface tension difference Δ*κ*_*s*_ indeed creates a recirculating flow pattern. This is because the surface tension gradient drives surface flows, a mechanism known as Marangoni effect [37]. For Δ*κ*_*s*_ *<* 0 the flow on the aggregate surface is oriented towards the anterior pole, similarly to what we observe in our experiments, while for Δ*κ*_*s*_ *>* 0 it is oriented towards the posterior pole (Figure 4d). This qualitative comparison suggests that in our gastruloids the surface tension of anterior regions may be higher than that of posterior regions.

We further quantitatively compared the flow fields obtained from our model to the experimentally observed one. We quantify the deviation between simulated and experimentally observed flow fields using a residual, which we define as the fraction of the spatial variance in the experimental flow field that is not reproduced by our simulations (see SI). In Figure 4f left, we plot the residual depending on the two model parameters Δ*κ*_*s*_ and *r*. We find indeed that the residual is smallest for negative Δ*κ*_*s*_. However, we also find that the deviation becomes larger for a small interface-to-surface stress ratio *r*. This is because, for small *r*, internally created tension are smaller than surface tension, which induces flows that would lead to a more circular aggregate shape (e.g. (Δ*κ*_*s*_, *r*) = (0, 0.5) in Figure 4d). However, we observe that even at negative Δ*κ*_*s*_ and large *r*, the residual is still very large, on the order of 85%.

To dissect the possible reasons for the relatively large residual between measured and predicted flow field (Figure 4f left), we suspected that small-scale spatio-temporal fluctuations contribute to the experimental flow fields. We thus compared between simulation and experiment only the flow contributions provided by the modes 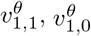 and 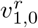, which together create the recirculating flow pattern. The residual between just these contributions between simulation and experiment is shown in Figure 4f right. We find that our simulations predict the observed recirculation flow in a wide parameter region defined by Δ*κ*_*s*_ *<* 0 with a residual that is reduced to ∼40 − 50%, independent of the precise values of Δ*κ*_*s*_ and *r*. However, varying Δ*κ*_*s*_ and *r* does affect the amplitude of other large-scale flow components that do not contribute to the recirculating flow. For example, reducing the values of *r* and |Δ*κ*_*s*_| results in an important flow contribution by the mode 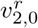 (Figure 4g), which corresponds to a rounding up of the gastruloid (e.g. (Δ*κ*_*s*_, *r*) = (0, 0.5) in Figure 4d).

Our results show that at the coarse-grained level, both the T/Bra and E-cad fields result in similar computational flow patterns (see also Figures S9, S10), as only the overall AP pattern is required to drive the main re-circulation flow. Taken together, the comparison between our model and experimental data suggests: (i) a higher surface tension in anterior (T/Bra −, E-cad−) than in posterior (T/Bra+, E-cad+) tissues and (ii) an interface tension between these tissues. To further test this, we performed fusion experiments between gastruloids.

### Fusion experiments confirm tissue-type-dependent tensions

To assess the existence of tissue-dependent tensions in the gastruloids, we performed fusion experiments to force posterior-like (T/Bra+) tissues to mechanically interact with anterior-like (T/Bra−) tissues (Figure 5, Supplementary Movie 7). To this end, we used our observations that gastruloids at 72hAAF display high T/Bra levels almost everywhere, while 84hAAF gastruloids are already polarized, exhibiting both T/Bra+ and T/Bra − regions. We thus brought 72hAAF gastruloids in contact with 84hAAF gastruloids.

**FIG. 5.**
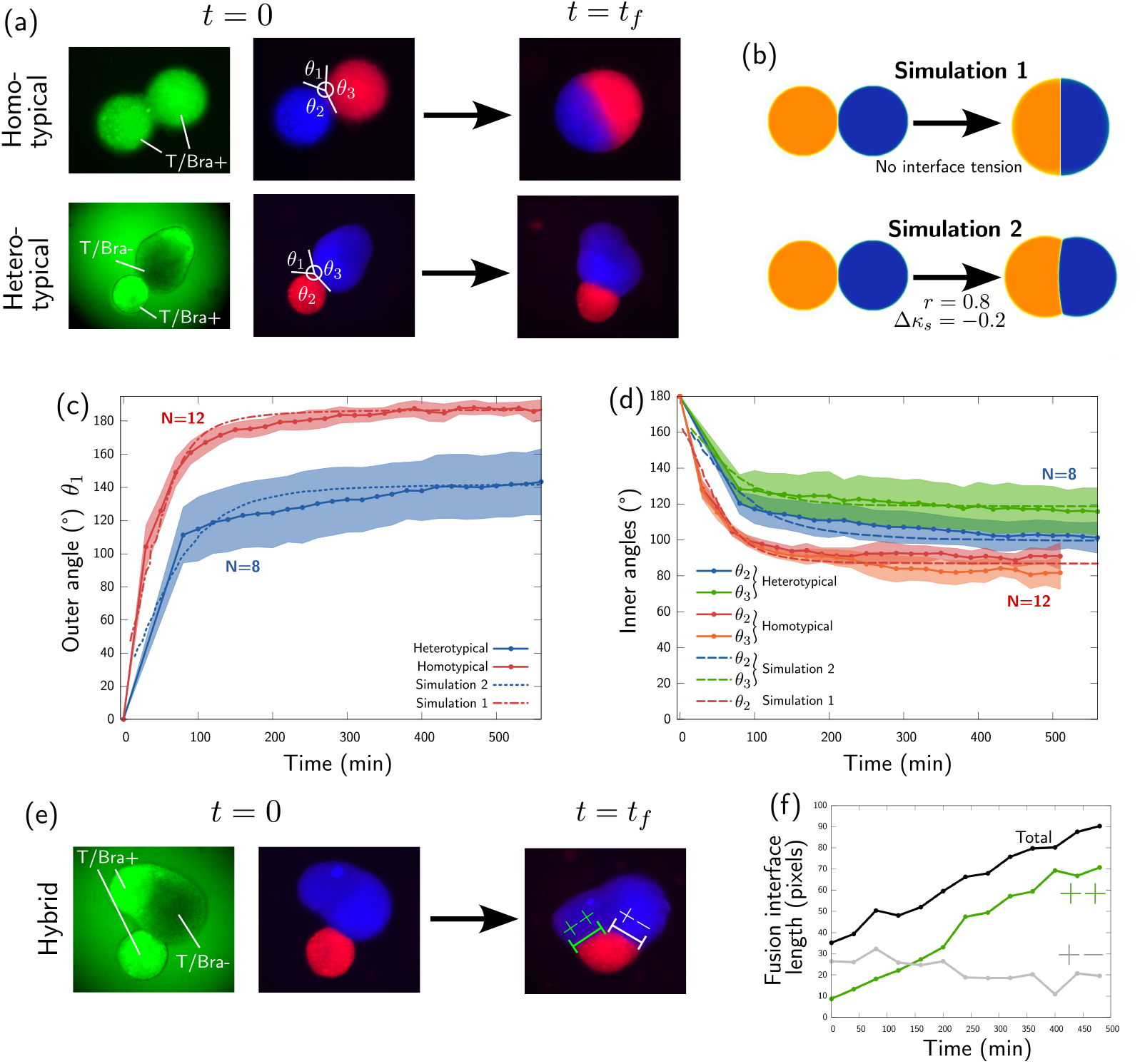
Quantitative analysis of aggregate fusion experiments. (a) Fusion experiments with either a homotypical ++ contact between two 72hAAF gastruloids or a heterotypical +− contact between a 72hAAF gastruloid and the T/Bra− region of a 84hAAF gastruloid. We define the fusion angles *θ*_1_, *θ*_2_ and *θ*_3_ as indicated. (b) Experiments are compared to model simulations of fusing droplets. we consider two particular cases: simulation 1 is performed in the absence of tension between the droplets, while simulation 2 is performed in the presence of interface tension and surface tension difference, namely *r* = 0.8 and Δ*κ*_*s*_ = − 0.2. (c,d) Time evolution of the (c) outer angle *θ*_1_ and (d) inner angles *θ*_2_ and *θ*_3_, averaged over samples and compared to the angles in the respective simulation. The shaded areas indicate the 95% confidence interval. (e) Example of hybrid fusion, where a 72hAAF gastruloid is in contact with both a T/Bra+ and a T/Bra− region of a 84hAAF gastruloid. (f) Time evolution of the fusion interface length, decomposed as the sum of homotypic contact length between two T/Bra+ tissue regions (++), and heterotypic contact length between a T/Bra+ and a T/Bra− region (+−).

Different fusion scenarios occurred depending on the relative position of both aggregates (Figure 5, Supplementary Movie 8). We first focus on interactions between T/Bra+ and T/Bra− tissues by selecting experiments with only a heterotypical + − contact of the 72hAAF gastruloid with a T/Bra− region of the 84hAAF gastruloid (Figure 5a). As a control experiment, we also consider homotypical ++ fusion experiments between two 72hAAF T/Bra+ aggregates (Supplementary Movie 9). We track the fusion angles *θ*_1_, *θ*_2_ and *θ*_3_ over time during the fusion process (Figure 5a,c,d, Methods). In the case of homotypical ++ fusions, the outer fusion angle *θ*_1_ saturates to 180^*°*^ (red solid curve in Figure 5c), while the inner angles *θ*_2_ and *θ*_3_ both saturate at approximately 90^*°*^ (red and orange solid curves in Figure 5d). This behavior is expected and quantitatively reproduced by our model when we set 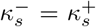 and *r* = 0 (Figure 5b, simulation 1; red dashed curves in Figure 5c,d).

For heterotypical + − fusions, the outer fusion angle saturates to a lower value of *θ*_1_ ≈ 140^*°*^ (blue solid curve in Figure 5c). Moreover, the inner angle *θ*_2_ in the T/Bra+ tissue saturates to a lower value than the inner angle *θ*_3_ in the T/Bra− tissue (blue and green solid curves, respectively, in Figure 5d). These observations are consistent with our previous conclusions that (i) T/Bra− tissues have a higher surface tension than T/Bra+ tissues (*θ*_2_ *< θ*_3_), and (ii) there is a finite interface tension between T/Bra+ and T/Bra tissues (*θ*_1_ *<* 180^*°*^). The latter observation is further supported by hybrid fusions, when a 72hAAF gastruloid was in contact with both T/Bra+ and T/Bra regions of a 84hAAF gastruloid (Figure 5e, see two cases in Supplementary Movie 8). In this case, we observed an asymmetric fusion, i.e. the homotypic interface between T/Bra+ and T/Bra+ regions (++) extends in length while the heterotypic interface between T/Bra+ and T/Bra regions (+) slightly shrinks (Figure 5f). This behavior suggests that the affinity between two T/Bra+ regions is larger than that between T/Bra+ and T/Bra regions. In other words, there is an effective interface tension between T/Bra+ and T/Bra− regions.

In our model, fusion angles are determined by tension ratios Δ*κ*_*s*_ and *r* (see SI). Simulations performed with Δ*κ*_*s*_ = − 0.2 and *r* = 0.8 quantitatively reproduce heterotypical fusion angles (Figure 5b, simulation 2; blue and green dashed curves in Figure 5c,d). As discussed before, in this part of parameter space, our simulations predict a recirculation flow together with a strong flow component relaxing gastruloids towards a circular shape (Figure 4g). We believe that in real gastruloids, such rounding up is counter-acted by additional mechanisms that lead to gastruloid elongation along the AP axis. However, an explicit tendency of the aggregates to elongate requires additional, active stresses, which are not included in our model, as we focus here on the recirculating flows and AP symmetry breaking.

While in the previous section the velocity amplitude was essentially a fit parameter (see SI), the fusion experiments allow us to independently predict the order of magnitude of the expected reciculating flow speed. The time scale on which two droplets with viscosity *η*, radius *R*, and surface tension *γ*_*s*_ fuse is given by *t*_*F*_ ∼ *Rη/γ*_*s*_ [38, 39]. According to Figure 5e, we estimate *t*_*F*_ ∼ 1 h. Moreover, in a droplet whose surface tension varies between *γ*_*s*−_ and *γ*^+^, recirculating Marangoni flows traverse the size of the droplet on the time scale 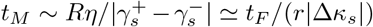 [40]. Using Δ*κ*_*s*_ = − 0.2 and *r* = 0.8, we thus obtain *t*_*M*_ ∼ 6 h, which is close to the time scale of gastruloid symmetry breaking *t*_*SB*_ ∼ 10 h. This confirms that the magnitude of the differential tension observed in our fusion experiments is sufficient to drive the observed recirculation flows, which promote symmetry breaking.

## III. DISCUSSION

In this paper, we study aggregates of mouse embryonic stem cells, which form a primary axis through rotational symmetry breaking, during which a pole of high T/Bra and E-Cad expression forms at one end (Figure 1). Several cellular mechanisms can contribute to such symmetry breaking (Figure 2c): biochemical reactions and cellular diffusion (RD), advection of regions with high T/Bra or E-Cad, and heterogeneous growth. We show that RD and advection both significantly contribute to polarization (Figure 2). RD contributions, important during the initial phases of symmetry breaking, are accompanied by significant advection contributions persisting over longer periods. We demonstrate that a recirculating component dominates these flows (Figure 3), and we show that the flows are consistent with Marangoni flows that are induced by differential tensions in an already polarized aggregate (Figure 4). We further confirm the existence of these tension differences by measuring interface angles in fusion experiments (Figure 5). Taken together, our observations suggest a positive feedback loop where recirculating flows amplify and thus stabilize an early aggregate polarization.

Since the middle of the 20th century, effective tissue surface tensions have been proposed to drive the separation of cell populations [31, 32, 41–45]. Such effective tensions can for instance be created by differential cadherin expression [32, 33]. Indeed, in our gastruloids, cells progressively lose E-cadherin expression [16], and display a spatial anti-correlation between N-cadherin and E-cadherin [16] and between OB-cadherin and E-cadherin (Supplementary Figure 16). However, much of the past work on differential tissue tensions driving the separation of cell populations considered tissue separation as a diffusive unmixing process, including in stem cell aggregates [46, 47]. In contrast, our work shows for the first time that differential tissue surface tensions can drive long-range hydrodynamic flows that are maintained over a long time (≳ 10 hours). In previous work, some of us have shown that such long-range flows can be important to substantially accelerate cellular unmixing [48].

There are a number of possible future extensions of this work. First, much of the past work on cellular unmixing focused on already differentiated cells [49]. However, if all cells kept their initial differentiation state over time, flows driven by differential tissue tensions may at some point cease [48]. Yet, the cells in our system are in the process of differentiating, and the observed recirculating flows are maintained over several hours, which makes us wonder whether the flows might be maintained through patterned cell differentiation. Indeed, in previous work, some of us have shown that (i) some T/Bra-negative cells present in the posterior pole are pluripotent cells, which divide and differentiate into T/Bra-positive cells, while (ii) the T/Bra+ and E-Cad-negative cells progressively divide and differentiate into T/Bra− and E-Cad-negative cells [16]. Whether and how precisely such cell division and differentiation events might help to sustain the observed flows will be interesting to dissect in future work.

Second, here we have studied a 2D section of the 3D flows in the aggregates. This is because of current technical limitations to *in toto* imaging of the aggregates, due to photo-toxicity constraints and line scanning speed limitations. Here, we have filtered out aggregates where the inferred out-of-plane polarization component was too high (see SI, Supplementary movie 10). It remains a future challenge to analyze the full 3D expression patterns and flows using *in toto* imaging of the aggregates.

Third, it will be interesting to examine the precise cellular mechanisms underlying the differences in effective tissue surface tension [31, 45, 50]. Possibilities are adhesion by itself, differential levels of cell contractility, or cell motility (Supplementary figures 16-17). These different microscopic contributions are not mutually exclusive and dissecting the exact contribution of each of them will necessitate the generation of specific knock-out mutant cell lines which is beyond the scope of this work.

Fourth, while we have focused here on the recirculating flows in the aggregate, we did not include any mechanism creating aggregate elongation, which would necessitate both active driving forces and globally aligned orientational information [51]. Indeed, it has been shown for fish embryonic stem cell aggregates, that cell polarity proteins are present and play an important role for gastruloid elongation [52, 53]. Alternative mechanisms could be based on heterogeneous mechanical properties [54]. It will be interesting to identify the precise mechanisms driving gastruloid elongation in future work.

We expect that Marangoni-effect-driven recirculating tissue flows operate also in many other organoid systems, i.e. systems formed from pluripotent cells. While such Marangoni flows can be artificially induced [55], they can also be driven internally. The emergence of such flows requires only the existence of distinct cell populations, which will typically display different surface tensions, thus providing a general mechanism for amplification of symmetry breaking. These flows may also be relevant in the gastrulating mouse embryo, where they interact with other internal and external mechanisms [2]. Specifically, during mouse gastrulation, epithelial cells of the epiblast undergo an EMT by downregulating adherens junction molecules such as E-cad. This allows the cells to delaminate from the epiblast and migrate away from the primitive streak as mesenchymal cells. There is evidence that these emerging mesenchymal cells migrate anteriorly as so-called embryonic mesodermal wings [56]. Moreover, recirculating flows have been identified during the formation of the first cardiac crescent, which takes place during gastrulation [3]. We thus anticipate that the feedback loop that we observe in our mouse embryonic stem cell aggregates may turn out to play a role in many systems both in vitro and in vivo.

## IV. METHODS

### Cell culture and gastruloid formation

We used the two cell lines, E-cad-GFP/Oct4-mCherry and T-Bra-GFP/NE-mKate2 (Histone GFP under the T-Bra promoter). In brief, between 100 and 200 cells were seeded in low adhesive 96 wells plates in a neural differentiation medium. In order to promote the formation of endoderm formation and to reduce symmetry breaking variability compared to the standard gastruloids protocol, we enrichted the medium with Fibroblast Growth Factor and Activin. A complete description of the cell lines, the culture conditions and the protocol for making gastruloids is presented in [16]. All cell lines were tested to be free from mycoplasma contamination using qPCR.

### Two-photon imaging

The gastruloids were imaged in a chamber maintained at 37^*°*^C, 5% CO2 with a humidifier. We performed two-photon imaging using a Zeiss 510 NLO (Inverse - LSM) with a femtosecond laser (Laser Mai Tai DeepSee HP) with a 40 x/1.2 C Apochromat objective. 72hAAF gastruloids were transferred from 96-well plates to either MatTek dishes (Mat-Tek corporation, ref: P35G-1.5-14 C) or to microwell plates (500 µm wells, SUNBIOSCIENCE ref: *Gri*3*D*), which enable to image in parallel up to four gastruloids thanks to the small distance between two neighbouring wells, see Supplementary Figure 18. To visualize inter-cellular space, we added 10 µL*/*mL of a Sulforhodamin B solution (1 mg*/*mL of Sulforhodamin B powder, 230,162 Aldrich) to the medium and excited both GFP and Sulforhodamin B at 900 nm. To image cell nuclei, we added 1 µL of a SPY555-DNA (spirochrome) 1000x stock solution per mL of medium and excited both GFP and SPY555-DNA at 880 nm. Nuclei staining progressively improved over time, likely due to a progressive decrease in cell packing density and to nuclei accessibility (Supplementary movies 2-5).

We imaged sub-volumes of gastruloids around their midplane (z-stacks of 60 µm, with 12 to 30 slices at intervals between 5 µm and 2 µm depending on the movie) for a total duration between 10 h and 24 h (time interval between two frames between 3 and 10 min depending on the experiment). We imaged the GFP and the red fluorophores (either Sulforhodamin B or SPY555-DNA) channels using two non-descanned detectors.

### Immunostaining

Samples were immunostained using the same protocol than in [16] apart from the fact that primary antibodies were incubated during 3 consecutive days. Secondary antibodies (AF488, A568, AF647) and Hoechst were chosen to be compatible with four-color two-photon imaging.

| Antibodies |  |  |  |  |
| --- | --- | --- | --- | --- |
| Antibody | Species | Reference | Provider | Dilution |
| Brachyury | Goat | AF2085 | RD system | 1:40 |
| E-cadherin | Rat | M108 | Takara | 1:200 |
| FoxA2 | Goat | sc-6554 | Santa-Cruz | 1:50 |
| Snail | Goat | AF3639 | RD system | 1:50 |
| OB-cadherin | Rabbit | 71-7600 | Invitrogen | 1:500 |
| Flk1 | Rat | 14-5821-85 | Ebioscience | 1:100 |
| VEGFR2 | Goat | AF644 | BioTechne | 1:200 |
| Snail | Rabbit | C15D3 | Cell Signaling | 1:100 |

### Optical flow measurements

To measure coarsegrained velocity fields, we use the signal collected in the red channel (either inter-cellular space with the Sulforhodamin B (SRB) or nuclear signal with the SPY555-DNA probe). The SRB signal corresponds to (1) cell contours, (2) dead cells, and (3) bright spots of subcellular size moving rapidly, corresponding probably to vesicles. As dead cells and bright spots are much brighter than cell contours, we filter them with a thresholding operation. The SPY555-DNA signal also accumulates more in bright spots and dead cells which are filtered by the same thresholding process. The mid-plane of the z-stack is determined (maximal area of the gastruloid) and an average projection is made between the mid-plane and the two planes under and above it (separated usually by 2 microns). The same averaging is performed on the GFP channel. We then perform 2D timelapse registration using the ImageJ Plugin *MultiStackReg* to remove rapid rotations and translations of the aggregate (the transformation is calculated on the red channel and applied to both red and green channels). We then use a custom-made Matlab optical flow code based on the Kanade Lucas Tomasi (KLT) algorithm with a level 2 pyramid and a window size corresponding to a square of 2 cells by 2 cells [28]. The velocity fields are averaged over 12 consecutive frames (1 hour) to smooth both measurement noise and temporal fluctuations. In Supplementary Figure 7, we compare the velocity maps obtained by applying the optical flow method on two distinct signals (SRB versus nuclei or raw nuclei signal versus locally equalized nuclei signal).

### Fusion experiments

Two gastruloid batches were prepared from the T/Bra GFP line with a 14 hours delay from one another. Both batches were incubated during the 24h long Chiron pulse in different CellTracker colors (CMTMR orange CellTracker, Thermofischer) for the older batch and blue (CMAC blue CellTracker, Thermofischer) for the youngest batch. They were rinsed in transparent medium (Neural differentiation medium enriched in FGF and Activin) at the time of fusion (t = 72hAAF for the youngest gastruloid and t = 86hAAF for the oldest one) and were put in contact in pairs (either a young/young pair or a young/old pair) in an individ-ual well from a 96 wells plate. Pairs of fusing gastruloids were imaged with epifluorescence and brightfield at 10x magnification with a Zeiss microscope every 20 minutes in the red, blue, green and brightfield channels using a multiposition configuration.

### Fusion interface/angle analysis

We used the T/Bra and CellTracker color channels to define three scalar fields: (i) a field *ψ*(***r***) equal to 1 inside any one of the two gastruloids and 0 outside, (ii) a field *χ*(***r***) equal to 1 in the first aggregate (e.g. blue in Figure 5c) and -1 in the second aggregate (e.g. red in Figure 5c), and (iii)a field *ϕ*(***r***) ∈ [0, 1] representing the T/Bra signal. We defined the fusion interface as the iso-line *χ* = 0 with curvilinear coordinate *s*_*χ*_. The fusion interface length was then defined as

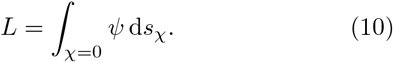

In addition, the length of the homotypic ++ interface between T/Bra+ tissues was computed as

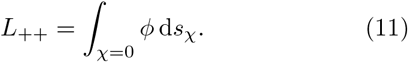

The heterotypic +− interface length was then computed as *L*_+−_ = *L* − *L*_++_.

We defined the gastruloids’ surface as the iso-line *ψ* = 0.5 with curvilinear coordinate *s*_*ψ*_. The triple points where both gastruloids and the outer medium meet are then characterized by the coordinate 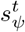 on the aggregate surface that satisfies 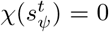. At a given triple point, we computed the outer fusion angle *θ*_1_ using

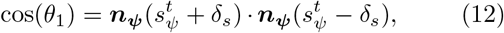

where ***n***_***ψ***_ =−∇*ψ/*|∇|*ψ* is the outward-pointing unit vector normal to the aggregate surface and *δ*_*s*_ is a length scale parameter. We checked that the choice of *δ*_*s*_ does not affect the results in Figure 5.

Inner angles *θ*_2_ and *θ*_3_ were determined following a similar approach, but using the gastruloid-gastruloid interface *χ* = 0 with curvilinear coordinate *s*_*χ*_. At a triple point, the coordinate 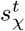 on the gastruloid-gastruloid interface satisfies 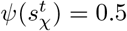. The inner angles were com-puted using

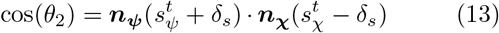

and

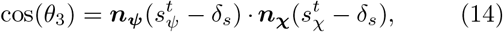

where ***n***_***χ***_ = ∇*χ/* |∇|*χ* is the unit vector normal to the fusion interface on the iso-line *χ* = 0.

## ACKNOWLEDGMENTS

M.M. thanks the Centre Interdisciplinaire de Nanoscience de Marseille (CINaM) for providing office space. This work is supported by the French National Research Agency (“France 2030”, ANR-16-CONV-0001 from Excellence Initiative of Aix-Marseille University - A*MIDEX and generic grant to P.-F.L. ANR-19-CE13-0022), the Fondation de la Recherche Médicale (to P.-F.L. EQU202003010407), a generic grant to S.T. ANR-22-CE30-0021 and an ERC grant (to M.M. and P.-F.L., ERC SyG 101072123). We also acknowledge the France-Bioimaging Infrastructure (ANR-10-INBS-04). S.T. thanks Daniel Sapede for his invaluable help on implementing multiposition timelapse.

## Footnotes

* This work is licensed under a Creative Commons “Attribution 4.0 International” license.

