## Supplementary Material for "Marangoni-like tissue flows enhance symmetry breaking of embryonic organoids"

#### 1 Dipole moment decomposition

##### Definition

We define the dipole moment of the 2D cross section $\Omega$ of the aggregate as

$$\boldsymbol{P}(t) = \int_\Omega \phi(\boldsymbol{r}, t) \big[\boldsymbol{r} - \boldsymbol{c}(t)\big] \, \mathrm{d}S. \tag{S1}$$

Here and below, $\boldsymbol{r}$ is the integration variable, $\phi(\boldsymbol{r}, t)$ is the protein fluorescence intensity (of T/Bra or E-cad), $\boldsymbol{c}(t) = \big(\int_\Omega \boldsymbol{r} \, \mathrm{d}S\big)/S(t)$ is the aggregate cross section's barycenter, and $S(t) = \int_\Omega \mathrm{d}S$ is its area.

##### Normalization

As a reference dipole, we consider a Janus particle, i.e. a disk with the same area $S$ as the aggregate cross section, and composed of two semi-circular surfaces with homogeneous concentrations $\tilde{\phi}$ and $-\tilde{\phi}$, respectively. The resulting reference dipole moment reads

$$P_0 = \frac{4}{3}\tilde{\phi}R^3, \tag{S2}$$

with $R = \sqrt{S/\pi}$ the radius of the Janus particle. For the characteristic concentration amplitude $\tilde{\phi}$, we choose the root-mean-square of the $\phi$-field within the aggregate cross section.

Since both $\tilde{\phi}$ and $R$ vary over time, we compared two different strategies for the normalization of the dipole moment:

1. Defining a time-independent $P_0$ according to Eq. (S2) based on $\tilde{\phi}$ and $R$ *at the end of the polarization process* (Figure S8b), or

2. defining a time-dependent $P_0(t)$ according to Eq. (S2) *using the instantaneous values of* $\tilde{\phi}(t)$ *and* $R(t)$ (Figure 2c in main text).

In the main text, we have decided to use the second strategy throughout, because we observed that it allows for a better collapse of the polarization curves of different aggregates (compare Figure 2c in main text to Figure S8b).

##### Decomposition

Using the Leibniz integral theorem, the time derivative of $\boldsymbol{P}$ reads

$$\frac{\mathrm{d}\boldsymbol{P}}{\mathrm{d}t} = \int_\Omega \frac{\partial}{\partial t}\big(\phi\,[\boldsymbol{r} - \boldsymbol{c}]\big)\,\mathrm{d}S + \int_{\partial\Omega} \phi\,[\boldsymbol{r} - \boldsymbol{c}](\boldsymbol{u}_b \cdot \boldsymbol{n}_{2D})\,\mathrm{d}l, \tag{S3}$$

where $\partial\Omega$ is the boundary of the aggregate cross section, $\boldsymbol{n}_{2D}$ is a unit vector within the cross-sectional plane normal to the boundary $\partial\Omega$, and $\boldsymbol{u}_b$ is the apparent boundary velocity in the cross-sectional plane. Since the aggregate can also move and deform in the direction normal to the cross-sectional plane, the apparent velocity $\boldsymbol{u}_b$ is the sum of (1) the components of the actual tissue velocity that lie in the cross-sectional plane, $\mathbf{v}_{2D}$, and (2) an apparent planar velocity resulting from motion of the boundary normal to the cross-sectional plane:

$$\boldsymbol{u}_b = \mathbf{v}_{2D} + \mathrm{v}_z n_z^* \boldsymbol{n}_{2D}, \tag{S4}$$

where $\mathrm{v}_z$ is the velocity normal to the cross-sectional plane, and $n_z^* = n_z/\sqrt{n_x^2 + n_y^2}$ with $\boldsymbol{n} = (n_x, n_y, n_z)$ being the 3D unit normal vector at the aggregate boundary.

Using the divergence theorem and denoting the gradient operator in the cross-sectional plane by $\nabla$, Eq. (S3) becomes

$$\frac{\mathrm{d}\boldsymbol{P}}{\mathrm{d}t} = \int_\Omega \frac{\partial}{\partial t}\left(\phi\,[\boldsymbol{r}-\boldsymbol{c}]\right)\mathrm{d}S + \int_\Omega \nabla\cdot\Big(\phi\mathbf{v}_{2D}[\boldsymbol{r}-\boldsymbol{c}]\Big)\,\mathrm{d}S + \boldsymbol{W}_{3D} \tag{S5}$$

with

$$\boldsymbol{W}_{3D} := \int_{\partial\Omega} \mathrm{v}_z n_z^* \phi\,[\boldsymbol{r}-\boldsymbol{c}]\,\mathrm{d}l. \tag{S6}$$

This can be written as

$$\frac{\mathrm{d}\boldsymbol{P}}{\mathrm{d}t} = \int_\Omega \left[\frac{\partial\phi}{\partial t} + \mathbf{v}_{2D}\cdot\nabla\phi\right][\boldsymbol{r}-\boldsymbol{c}]\;\mathrm{d}S - \frac{\mathrm{d}\boldsymbol{c}}{\mathrm{d}t}S\overline{\phi} + \int_\Omega \phi\Big(\mathbf{v}_{2D} + (\nabla\cdot\mathbf{v}_{2D})[\boldsymbol{r}-\boldsymbol{c}]\Big)\,\mathrm{d}S + \boldsymbol{W}_{3D}, \tag{S7}$$

where $\overline{\phi} := (\int_\Omega \phi\,\mathrm{d}S)/S$. The first term on the right-hand side of Eq. (S7) represents the reacton-diffusion contributions to dipole moment variations, the second and third correspond to advective contributions, and the last term, $\boldsymbol{W}_{3D}$, represents 3D effects.

To further reduce the 3D effect contribution, we apply the Leibniz rule to $\mathrm{d}\boldsymbol{c}/\mathrm{d}t$ in Eq. (S7):

$$\frac{\mathrm{d}\boldsymbol{c}}{\mathrm{d}t} = -\frac{\boldsymbol{c}}{S}\frac{\mathrm{d}S}{\mathrm{d}t} + \frac{1}{S}\int_\Omega(\nabla\cdot\mathbf{v}_{2D})\boldsymbol{r}\,\mathrm{d}S + \overline{\mathbf{v}_{2D}} + \frac{1}{S}\int_{\partial\Omega}\mathrm{v}_z n_z^* \boldsymbol{r}\,\mathrm{d}l, \tag{S8}$$

where $\overline{\mathbf{v}_{2D}} := (\int_\Omega \mathbf{v}_{2D}\,\mathrm{d}S)/S$. Substituting Eq. (S8) into Eq. (S7), we obtain

$$\frac{\mathrm{d}\boldsymbol{P}}{\mathrm{d}t} = \boldsymbol{R} + \boldsymbol{A} + \boldsymbol{G} + \boldsymbol{Q}_{3D}, \tag{S9}$$

with

$$\boldsymbol{R}(t) := \int_\Omega \left[\frac{\partial\phi}{\partial t} + \mathbf{v}_{2D}\cdot\nabla\phi\right][\boldsymbol{r}-\boldsymbol{c}]\;\mathrm{d}S \qquad \text{(reaction-diffusion)} \tag{S10}$$

$$\boldsymbol{A}(t) := \int_\Omega \phi\,[\mathbf{v}_{2D} - \overline{\mathbf{v}_{2D}}]\,\mathrm{d}S \qquad \text{(advection)} \tag{S11}$$

$$\boldsymbol{G}(t) := \int_\Omega (\phi - \overline{\phi})(\nabla\cdot\mathbf{v}_{2D})[\boldsymbol{r}-\boldsymbol{c}]\,\mathrm{d}S, \qquad \text{(growth)} \tag{S12}$$

and where we introduced the new 3D correction:

$$\boldsymbol{Q}_{3D}(t) := \int_{\partial\Omega} \mathrm{v}_z n_z^* (\phi - \overline{\phi})\,[\boldsymbol{r}-\boldsymbol{c}]\,\mathrm{d}l. \tag{S13}$$

The 3D effects contribution $\boldsymbol{Q}_{3D}$ is generally expected to be smaller than $\boldsymbol{W}_{3D}$ defined in Eq. (S6), because the definition of $\boldsymbol{Q}_{3D}$ is reduced by the "mean-field 3D effects $\boldsymbol{W}_{3D}$" that assume a homogeneous

$\phi$ field. Therefore, in the paper we use Eq. (S9) to quantify contributions to dipole moment variations in the cross-sectional plane, where for notational simplicity $\mathbf{v}_{2D}$ is replaced by $\mathbf{v}$.

##### Analysis procedure

We use our live imaging data to quantify each of the terms in Eq. (S9). The velocity field $\mathbf{v}_{2D}$ is obtained from an optical flow analysis (Figure 1c left in the main text, Figure S6), and the scalar field $\phi$ corresponds to T/Bra or Ecad fluorescence intensity (Figure 1c right in the main text, Figure S1 and Figure S4). We study the dynamics of 10 gastruloids, including 6 for the T/Bra reporter line and 4 for the E-cad reporter line. The cumulative polarization change during the acquisition time interval $[t_i, t_f]$ is

$$\tilde{\boldsymbol{P}}(t) := \int_{t_i}^{t} \frac{\mathrm{d}\boldsymbol{P}}{\mathrm{d}t} dt = \tilde{\boldsymbol{R}}(t) + \tilde{\boldsymbol{A}}(t) + \tilde{\boldsymbol{G}}(t) + \tilde{\boldsymbol{Q}}_{3D}(t), \tag{S14}$$

where $\tilde{\boldsymbol{R}}$, $\tilde{\boldsymbol{A}}$, and $\tilde{\boldsymbol{G}}$ are computed as the time integrals of Eqs. (S10), (S11), and (S12), respectively. Because we cannot extract the precise values of $\mathrm{v}_z n_z^*$ along the boundary, we determined the 3D contribution $\tilde{\boldsymbol{Q}}_{3D}$ as

$$\tilde{\boldsymbol{Q}}_{3D}(t) = \tilde{\boldsymbol{P}}(t) - \tilde{\boldsymbol{R}}(t) - \tilde{\boldsymbol{A}}(t) - \tilde{\boldsymbol{G}}(t), \tag{S15}$$

where $\tilde{\boldsymbol{P}}(t)$ is computed based on equation (S1).

We discarded samples that showed too strong 3D effects; more precisely, we discarded the samples where $|\tilde{\boldsymbol{Q}}_{3D}(t_f)|/|\tilde{\boldsymbol{P}}(t_f)| > 0.2$ at the end of the acquisition time, $t_f$. The decomposition of the cumulative polarization change for the 5 resulting samples is plotted in Figure S8c-e,g,i. Two of the five discarded samples are shown in Figure S8f,h.

#### 2 Mode decomposition of tissue flows

##### Flow decomposition

In a disk of radius $R$, we express tissue velocity in polar coordinates $\mathbf{v}(r,\theta) = [\mathrm{v}_r(r,\theta), \mathrm{v}_\theta(r,\theta)]$, which we decompose into base modes $v^r_{n,p}(r,\theta)$ and $v^\theta_{n,p}(r,\theta)$ as follows:

$$\mathrm{v}_r(r,\theta) = \sum_{n=0}^{+\infty}\sum_{p=0}^{+\infty} A^r_{n,p} v^r_{n,p}(r,\theta), \tag{S16a}$$

$$\mathrm{v}_\theta(r,\theta) = \sum_{n=0}^{+\infty}\sum_{p=0}^{+\infty} A^\theta_{n,p} v^\theta_{n,p}(r,\theta), \tag{S16b}$$

where $A^r_{n,p}$ and $A^\theta_{n,p}$ are complex amplitudes.

We construct the base modes from a radial part given by polynomials with real coefficients, $\mathcal{P}_p(r)$, and an angular part given by Fourier modes with index $n$:

$$v^r_{n,p}(r,\theta) = v^\theta_{n,p}(r,\theta) = \mathcal{P}_p(r)\frac{e^{in\theta}}{\sqrt{2\pi}}. \tag{S17}$$

Thus, both radial and angular modes, $v^r_{n,p}(r,\theta)$ and $v^\theta_{n,p}(r,\theta)$, are formally defined in the same way. We still use a different notation only to differentiate whether they are used to construct the radial or the angular part of the velocity field $\mathbf{v}(\boldsymbol{r})$.

For convenience, we want these base modes to fulfill the following orthonormality condition:

$$\int_0^R \int_0^{2\pi} v^r_{n,p}(r,\theta)\left[v^r_{n',p'}(r,\theta)\right]^* r\, \mathrm{d}\theta \mathrm{d}r = \delta_{nn'}\delta_{pp'}, \tag{S18}$$

where the asterisk * denotes complex conjugation. This implies that

$$\int_0^R r\mathcal{P}_p(r)\mathcal{P}_{p'}(r)\,\mathrm{d}r = \delta_{pp'}. \tag{S19}$$

We thus constructed the polynomials $\mathcal{P}_p(r)$ following a standard modified Gram–Schmidt (MGS) procedure with the following inner product

$$\langle f, g\rangle = \int_0^R r f(r)g(r)\,\mathrm{d}r, \tag{S20}$$

with $f$ and $g$ two continuous functions on $[0, R]$. This procedure leads to a variant of the so-called Jacobi polynomials and ensures validity of Eqs. (S18) and (S19). In our construction, we ensured that each polynomial $\mathcal{P}_p(r)$ is of degree $p$, respectively (compare Figure 3c in the main text).

As a consequence of having a complete orthonormal basis, we can extract the mode amplitudes $A^r_{n,p}$ and $A^\theta_{n,p}$ as follows:

$$A^r_{n,p} = \int_0^R \int_0^{2\pi} \mathrm{v}_r(r,\theta)\left[v^r_{n,p}(r,\theta)\right]^* r\,\mathrm{d}\theta\mathrm{d}r, \tag{S21a}$$

$$A^\theta_{n,p} = \int_0^R \int_0^{2\pi} \mathrm{v}_\theta(r,\theta)\left[v^\theta_{n,p}(r,\theta)\right]^* r\,\mathrm{d}\theta\mathrm{d}r. \tag{S21b}$$

Moreover, the overall variance of the flow field can be expressed as follows (Parseval's identity):

$$V := \int_C \mathbf{v}^2(\boldsymbol{r})\,\mathrm{d}A = \sum_{n=0}^{+\infty}\sum_{p=0}^{+\infty}\left(|A^r_{n,p}|^2 + |A^\theta_{n,p}|^2\right). \tag{S22}$$

Here, $C$ denotes the circular disk of radius $R$. Thus, the spatial variance of the velocity field (up to the area factor) corresponds to the sum of the absolute squares of the complex mode amplitudes. This allows us to quantify the contribution of a given mode, e.g. $v^r_{n,p}$, to the overall spatial flow field fluctuations, which is given by $|A^r_{n,p}|^2/V$. Correspondingly, in Figures 3f,g and 4g in the main text, we normalized the squared amplitudes $|A^r_{n,p}|^2$ and $|A^\theta_{n,p}|^2$ by $V$.

**Recirculation mode**

We define the recirculation mode as

$$\mathbf{v}_R(r,\theta) = A^r_{1,0}v^r_{1,0}\boldsymbol{e}_r + \left[A^\theta_{1,1}v^\theta_{1,1} + A^\theta_{1,0}v^\theta_{1,0}\right]\boldsymbol{e}_\theta, \tag{S23}$$

where the complex amplitudes $A^r_{1,0}$, $A^\theta_{1,1}$ and $A^\theta_{1,0}$ are determined from the measured flow field $\mathbf{v}(\boldsymbol{r})$ through Eqs. (S21). In order to determine a statistical representation of the recirculation mode over samples, we compute these complex amplitudes for all samples and then define median amplitudes $\overline{A^r_{1,0}}$, $\overline{A^\theta_{1,1}}$ and $\overline{A^\theta_{1,0}}$, obtained by taking the median of the norms and the median of the phases. In Figure 3i in the main text, we use the phase of $\overline{A^r_{1,0}}$ to define the orientation of the recirculation mode.

### 3 Computational model

**Phase field model**

We model gastruloids as incompressible viscous fluids using Eqs. (5) and (6) in the main text. The internally generated stress $\boldsymbol{\sigma}_i$ is computed based on $\phi$ as in Eq. (6) in the main text with coefficient $\kappa_i$, and the surface stress $\boldsymbol{\sigma}_s$ is computed analogously, replacing $\phi$ by $\psi$ and $\kappa_i$ by $\kappa_s$. However, the aggregate surface stress also depends on the field $\phi(\boldsymbol{r})$ representing T/Bra or E-cad concentration. To account for this $\phi$-dependence of the surface stress, we use a single free-energy functional defining the thermodynamics of the ternary mixture composed of T/Bra+ tissues, T/Bra− tissues and external medium, from which the driving force can be derived as discussed in [2, 3]. The effective stresses only depend on 3 parameters controlling the respective interface tensions between T/Bra+ and T/Bra− tissues ($\kappa_i$), the surface tension between T/Bra+ tissues and the medium ($\kappa_s^+$) and the surface tension between T/Bra− tissues and the medium ($\kappa_s^-$).

##### Computational approach

We use experimental T/Bra fields to compute the stresses $\boldsymbol{\sigma}_i$ and $\boldsymbol{\sigma}_s$. The tissue velocity $\mathbf{v}$ and pressure $\Pi$ are then computed using a lattice-Boltzmann method [3, 4] accelerated by a multigrid method [5]. The computational procedure can be summarized as follows:

- We mapped the T/Bra fluorescence signal in the cross-sectional plane to the phase field $\phi$ by rescaling its magnitude by a global factor and adding a global offset, such that the maximum intensity is at $\phi = 1$ and the minimum at $\phi = -1$. We further defined the phase field $\psi$ equal to 1 inside the aggregate and 0 outside, which we also did using the T/Bra channel.
- We smoothed both $\phi$ and $\psi$ using convolutions with a Gaussian with a standard deviation of $\sim 20\,\mu$m.
- We projected the $\phi$ and $\psi$ fields onto a $256 \times 256$ grid that is suitable for computational simulations. In both the $x$ and $y$ directions, the grid extends over approximately twice the gastruloid size, allowing a suitable resolution of the interface between the gastruloid and the outer medium.
- Based on the stress coefficients $\kappa_i$, $\kappa_s^+$ and $\kappa_s^-$, we compute the force driving tissue flows using the approach from [2, 3].
- We perform lattice-Boltzmann iterations until a steady flow solution is obtained.

An overview of the computational results for all samples is provided in Figure S9 and Figure S10.

##### Computational residual

We perform simulations over ranges of the stress coefficients $\kappa_s^+$, $\kappa_s^-$ and $\kappa_i$. In particular, tissue flows depend on two non-dimensional ratios,

$$r = \frac{2\kappa_i}{\kappa_s^+ + \kappa_s^-} \tag{S24}$$

and

$$\Delta\kappa_s = \frac{\kappa_s^+ - \kappa_s^-}{\kappa_i}. \tag{S25}$$

A third parameter $\kappa_i/\eta$ sets the tissue velocity magnitude, but does not affect the flow pattern. We thus define the computational residual as

$$R(c) = \frac{\langle(\boldsymbol{u} - c\mathbf{v})^2\rangle}{\langle\boldsymbol{u}^2\rangle}, \tag{S26}$$

where $\langle\cdot\rangle$ corresponds to an average over the whole aggregate cross-section, $\boldsymbol{u}$ and $\mathbf{v}$ are the experimental and computational residuals, respectively, and $c$ is a free coefficient scaling the computational velocity magnitude. We can show analytically that the residual is minimal for $c = \langle \boldsymbol{u} \cdot \mathbf{v} \rangle / \langle \mathbf{v}^2 \rangle$. The minimal residual thus reads

$$R = 1 - \frac{\langle \boldsymbol{u} \cdot \mathbf{v} \rangle^2}{\langle \boldsymbol{u}^2 \rangle \langle \mathbf{v}^2 \rangle}. \tag{S27}$$

In the paper, we used Eq. (S27) to compute the computational residual (Figure 4f in the main text).

### 4 Contact angles between T/Bra+ and T/Bra− tissues

Our mechanical model assumes the existence of an interface tension between T/Bra+ and T/Bra− tissues and a surface tension between these tissues and the outer medium. In this framework, T/Bra+ and T/Bra- tissues thus behave as two immiscible fluid droplets and we can thus use classical theory for wetting mechanics to predict contact angles between both tissues at mechanical equilibrium.

We consider the ternary fluid configuration represented in Figure S13. Interfaces between fluids 1, 2 and 3 are governed by surface tensions $\sigma_{12}$, $\sigma_{13}$ and $\sigma_{23}$. At mechanical equilibrium, the balance between tension forces at one triple point results in the Neumann triangle relation,

$$\frac{\sigma_{12}}{\sin(\theta_3)} = \frac{\sigma_{23}}{\sin(\theta_1)} = \frac{\sigma_{13}}{\sin(\theta_2)}. \tag{S28}$$

As a result, the equilibrium angles $\theta_1$, $\theta_2$ and $\theta_3$ are entirely determined by the surface tensions $\sigma_{12}$, $\sigma_{13}$ and $\sigma_{23}$.

The full geometry of the equilibrium configuration can be computed considering the conserved area of droplets 2 and 3. At equilibrium, fluid 2 forms a lens characterized by one arc of radius $R_2$ and angle $\alpha_{21}$ and one arc of radius $R_{23}$ and angle $\alpha_{23}$. Similarly, fluid 3 forms a lens characterized by two arcs of radii $R_3$ and $R_{23}$ and angles $\alpha_{31}$ and $\alpha_{23}$. Angles $\theta_1$, $\theta_2$, $\theta_3$ and $\alpha_{21}$, $\alpha_{31}$, $\alpha_{23}$ are related through

$$\theta_3 = \frac{1}{2}(\alpha_{31} - \alpha_{23}), \tag{S29}$$

$$\theta_2 = \frac{1}{2}(\alpha_{21} - (2\pi - \alpha_{31})) \tag{S30}$$

and $\theta_1 = 2\pi - \theta_2 - \theta_3$.

The area $A_2$ of fluid 2 can be expressed as

$$A_2 = R_2^2 \left[ \alpha_{21} - \pi - \frac{1}{2}\sin(\alpha_{21}) \right] + R_{23}^2 \left[ \alpha_{23} - \frac{1}{2}\sin(\alpha_{23}) \right]. \tag{S31}$$

Similarly, the area of fluid 3 is

$$A_3 = R_3^2 \left[ \alpha_{31} - \pi - \frac{1}{2}\sin(\alpha_{31}) \right] - R_{23}^2 \left[ \alpha_{23} - \frac{1}{2}\sin(\alpha_{23}) \right]. \tag{S32}$$

Together with the extra condition required to ensure the connectivity of the three arc lengths, equations (S31) and (S32) allow to determine the unknown radii $R_2$, $R_3$ and $R_{23}$ from the conserved fluid area $A_2$ and $A_3$.

### 5 Supplementary movies

##### Supplementary movie 1

One hour long timelapse of 3 gastruloids used to compute recirculation velocity fields presented in Supplementary Figure 5 (top 78 hAAF, middle 80 hAAF and bottom 86 hAAF). On the left, T/Bra signal and on the right Sulforhodamin B signal. Both signals are inverted for better visibility.

**Supplementary movie 2**

z-stack of a 86 hAAF T/Bra GFP gastruloid. On the left in green, T/Bra signal and on the right in purple, SPY555-DNA signal.

**Supplementary movie 3**

Timelapse of a T/Bra GFP gastruloid between 76 hAAF and 91 hAAF. On the left in green, T/Bra signal and on the right in purple, SPY555-DNA signal.

**Supplementary movie 4**

z-stack of a 76 hAAF E-cadherin GFP gastruloid. On the left in green, E-cadherin signal and on the right in purple, SPY555-DNA signal.

**Supplementary movie 5**

Timelapse of a E-cadherin GFP gastruloid between 76 hAAF and 97 hAAF. On the left in green, E-cadherin signal and on the right in purple, SPY555-DNA signal.

**Supplementary movie 6**

Timelapse of a recirculation velocity field (moving average during one hour) corresponding to the Supplementary movie 3.

**Supplementary movie 7**

3 hours long epifluorescence timelapse of fusions between 72 hAAF gastruloids (in blue Cell Tracker) with 84 hAAF gastruloids (in orange Cell Tracker).

**Supplementary movie 8**

Zoom on few representative cases of different fusion scenarios that depend on the relative position of both aggregates extracted from Supplementary movie 8.

**Supplementary movie 9**

16 hours long epifluorescence timelapses of homogeneous fusions between 72 hAAF gastruloids (in blue Cell Tracker) and 72 hAAF gastruloids (in orange Cell Tracker).

**Supplementary movie 10**

Example of timelapse that has been discared because the inferred out-of-plane flow component was too high.

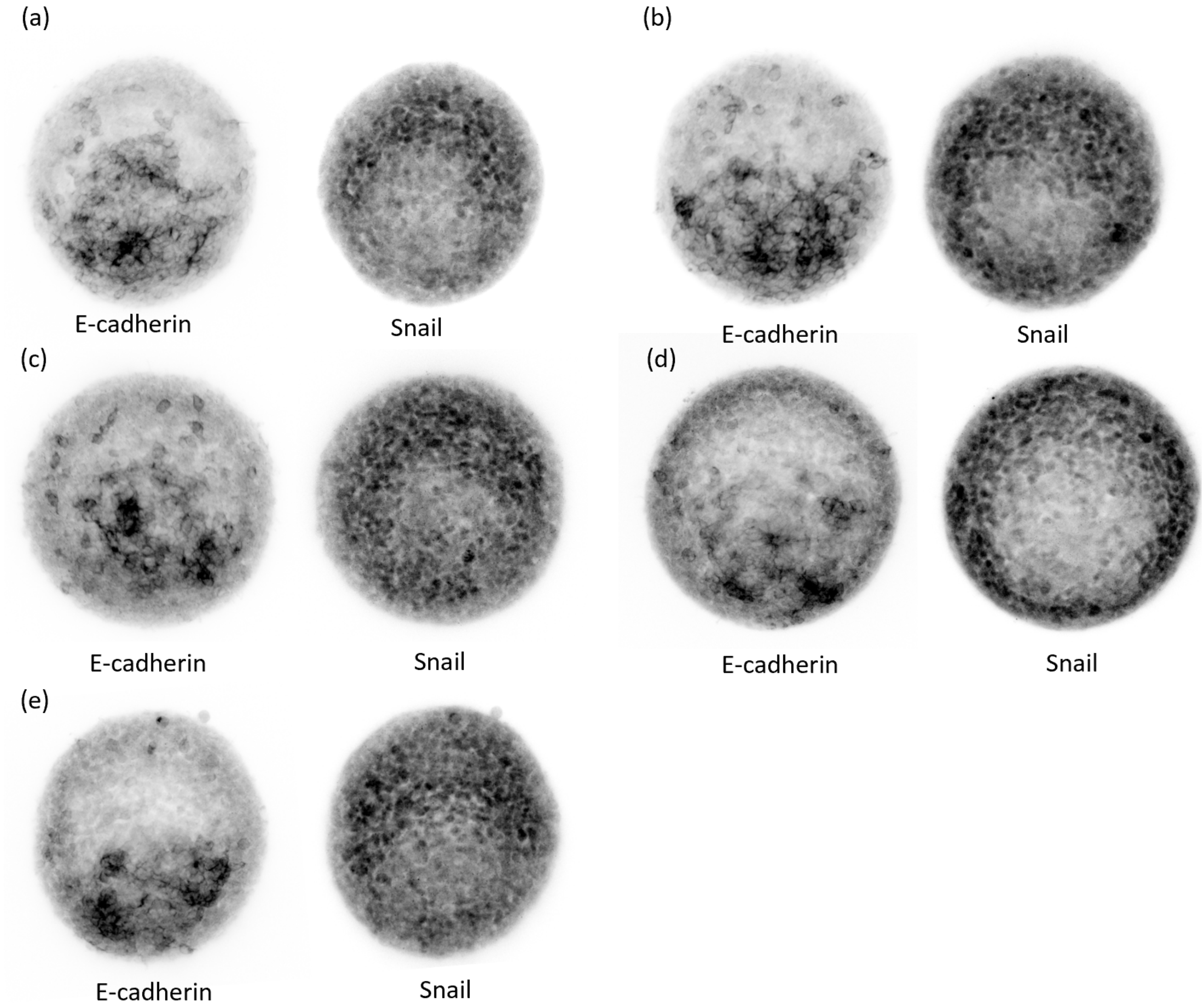


Figure S1: Epithelial-mesenchymal transition (EMT). (a-e) Immunostaining of E-cadherin and Snail for 5 different 80hAAF gastruloids (see Methods, Immunostainings). Each image corresponds to an average projection of a 20 µm-thick stack (20 slices), centered around the gastruloid mid-plane. Images are inverted (high-intensity pixels appear in black) for better visibility.

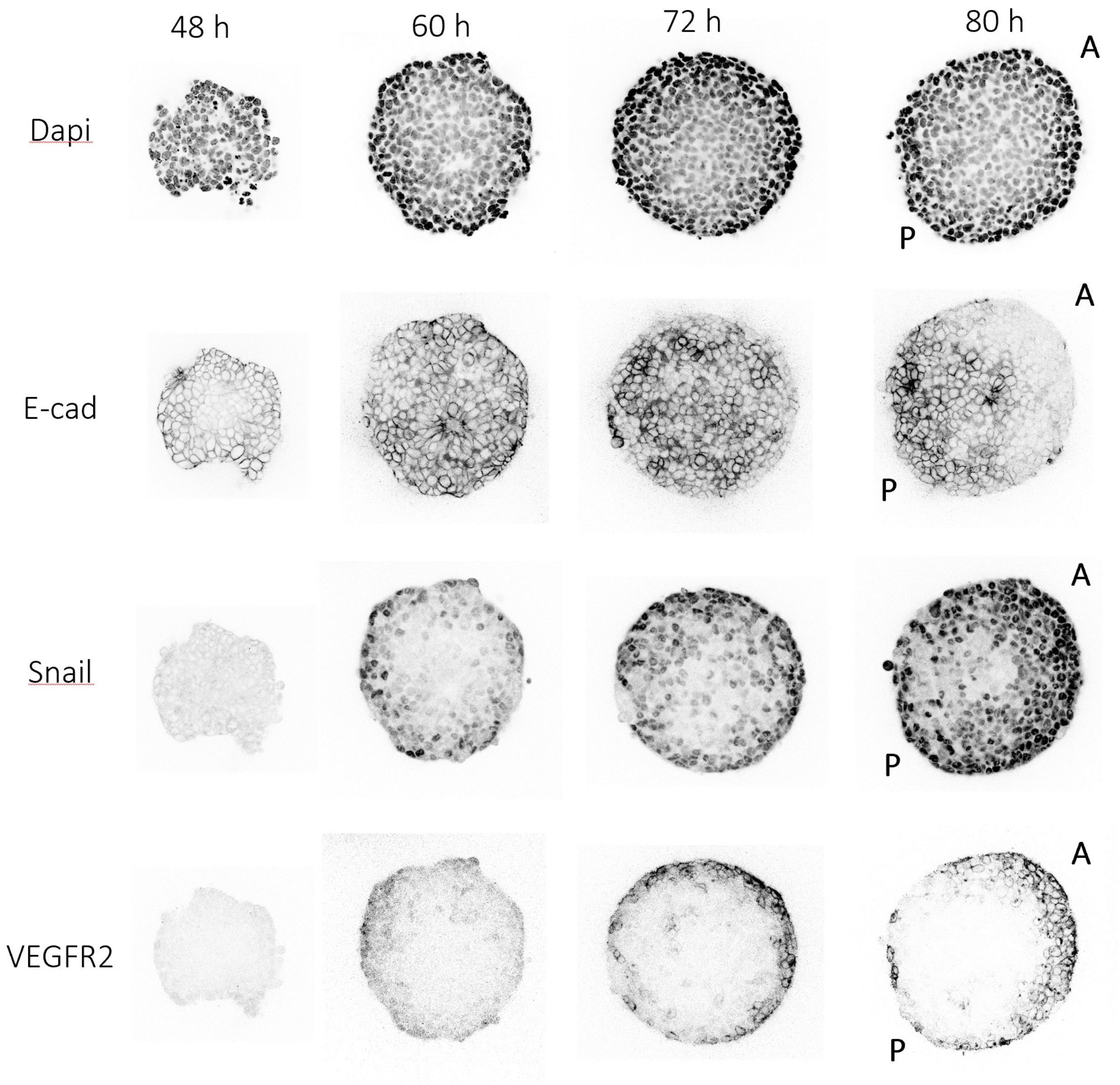


Figure S2: Epithelial-mesenchymal transition evolution before and during symmetry breaking. Immunostainings of gastruloids between 48 hAAF and 80 hAAF with E-Cadherin, Snail, VEGFR2 and Dapi labeled (see Methods, Immunostainings). At 48 hAAF (just before the Chiron pulse), cells are all E-Cadherin positive and both Snail and Vegfr2 negative. At the middle of the 24h-long Chiron pulse (60 hAAF), Snail is expressed by a large proportion of cells and cells at the periphery of the gastruloid start to lose their E-Cadherin expression. At the end of the Chiron pulse (72 hAAF), a higher proportion of cells express Snail near the gastruloid surface and are both E-Cadherin negative and VEGFR2 positive. The center of the gastruloid is composed of a globally E-cadherin positive population with sparse cells which are Snail positive and E-cadherin negative. At 80 hAAF, the E-Cadherin negative and VEGFR2 positive cell population appears both at the anterior part of the gastruloid and at the surface of its posterior part. The letter A denotes for the anterior pole and P for the posterior one.

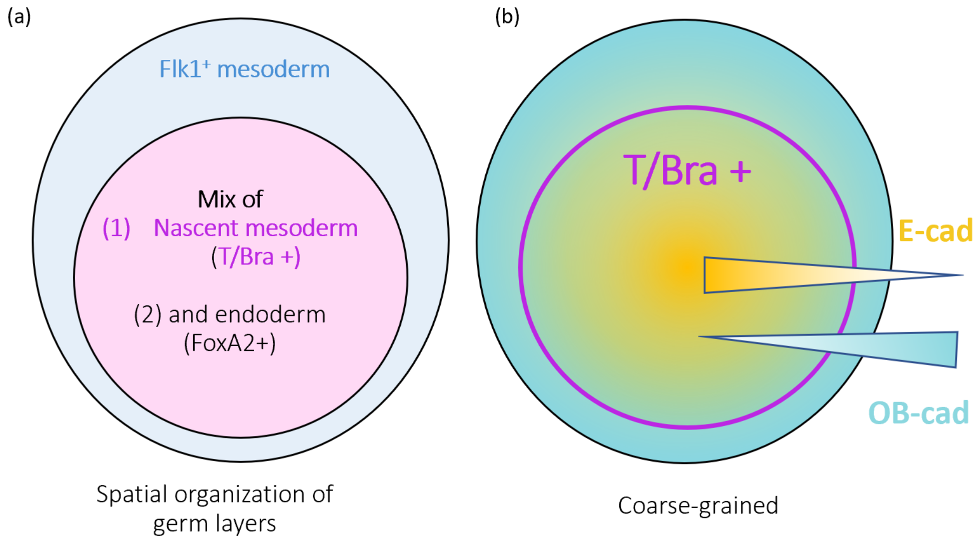


Figure S3: Schematics illustrating (a) the different cell populations within our gastruloids and (b) differential cadherin expression within these populations. (a) At 80hAAF, the gastruloids exhibit a stereotypical spatial separation between different cell populations. One pole expresses T/Bra, which indicates nascent mesoderm, mixed in a salt-and-pepper manner with cells expressing FoxA2, which indicates endoderm. The complementary pole is characterized by the absence of T/Bra and FoxA2 positive cells and the expression of Flk-1, which indicates more differentiated mesoderm, since Flk-1 is a hemato-cardiovascular cell lineages marker [1]. (b) At the coarse-grained level, the pole with many T/Bra-positive cells expresses high levels of E-cadherin and low levels of OB-cadherin, while the complementary population (Flk1 positive) expresses high levels of OB-cadherin and low levels of E-cadherin.

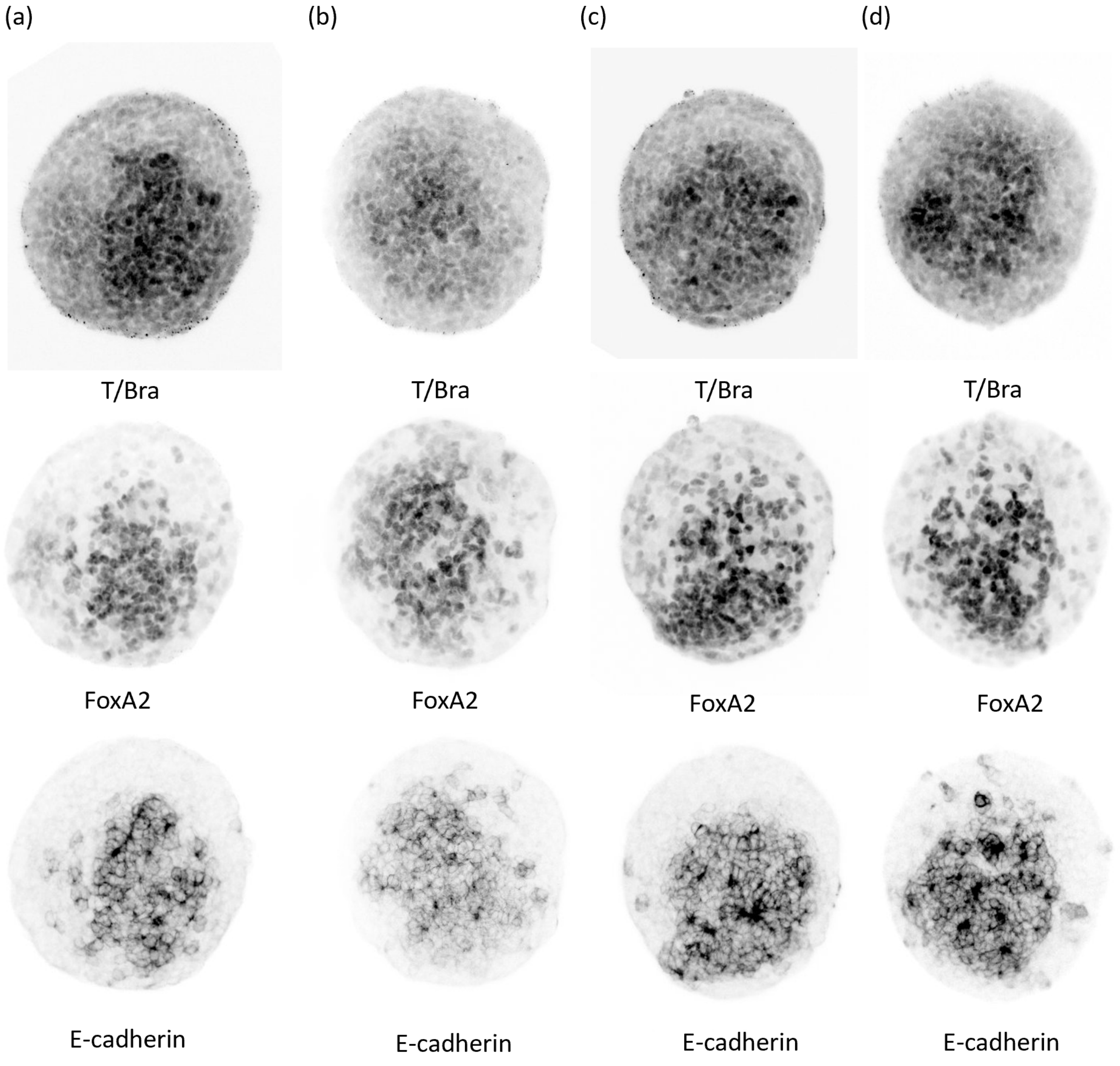


Figure S4: The E-cadherin positive population is a mix of endoderm (FoxA2+ cells) and nascent mesoderm (T/Bra+ cells). (a-d) Immunostaining of T/Bra, FoxA2 and E-cadherin for 4 different 80hAAF gastruloids (see Methods, Immunostainings). Each image corresponds to an average projection of a 20 µm-thick stack (20 slices) centered around the gastruloid mid-plane. Images are inverted (high-intensity pixels appear in black) for better visibility.

(a)

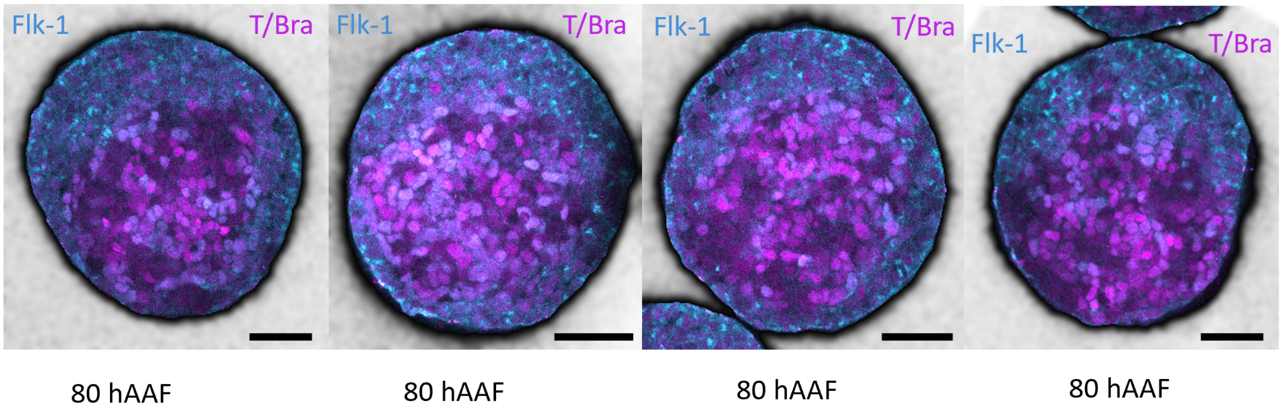


Figure S5: (a) Immunostaining of T/Bra and Flk-1 for 4 different 80hAAF gastruloids (see Methods, Immunostainings).

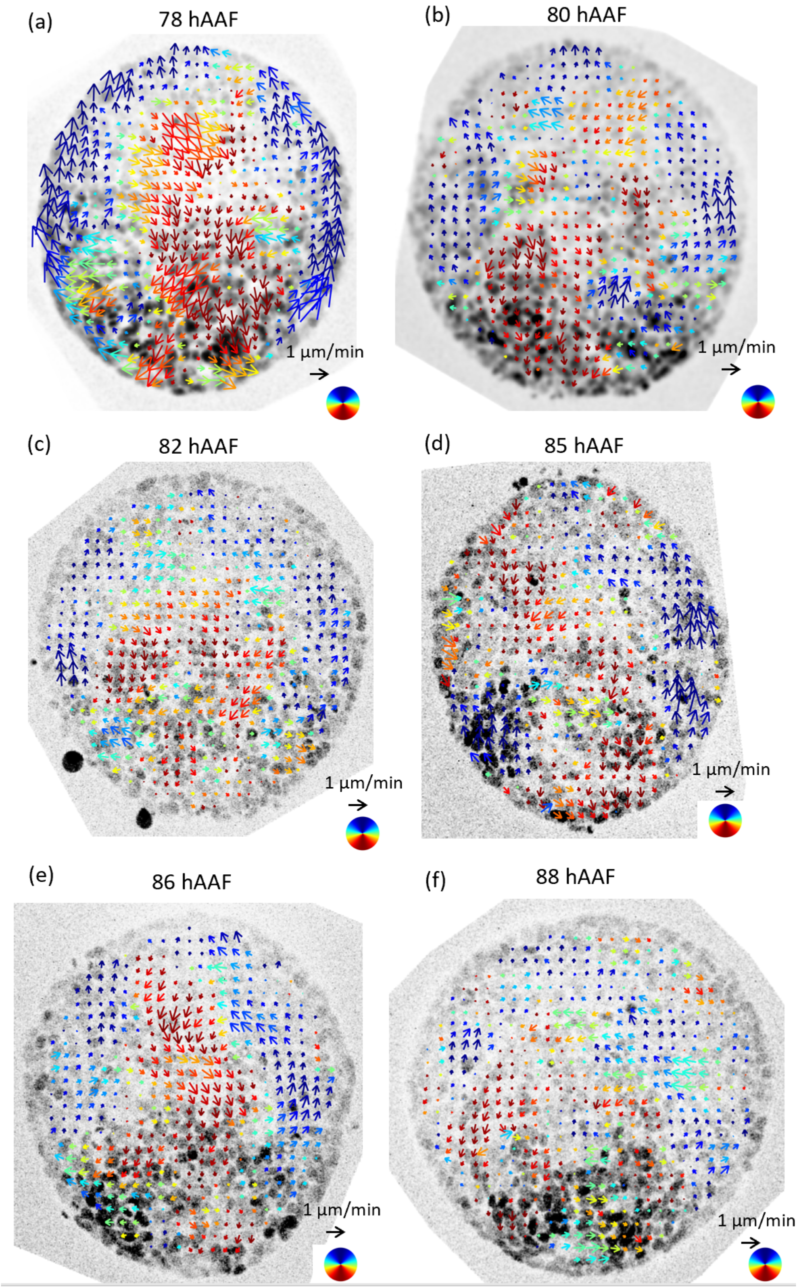


Figure S6: Quantified velocity fields associated to the symmetry breaking superimposed on the T/Bra field (inverted for better visibility). (a-f) Velocity fields are obtained by averaging over one hour (12 consecutive frames) the velocity fields were obtained by optical flow, performed on an average projection of a 10 μm-thick stack (5 slices, Sulforhodamin B) centered around the gastruloid mid-plane (see Methods, Optic Flow).

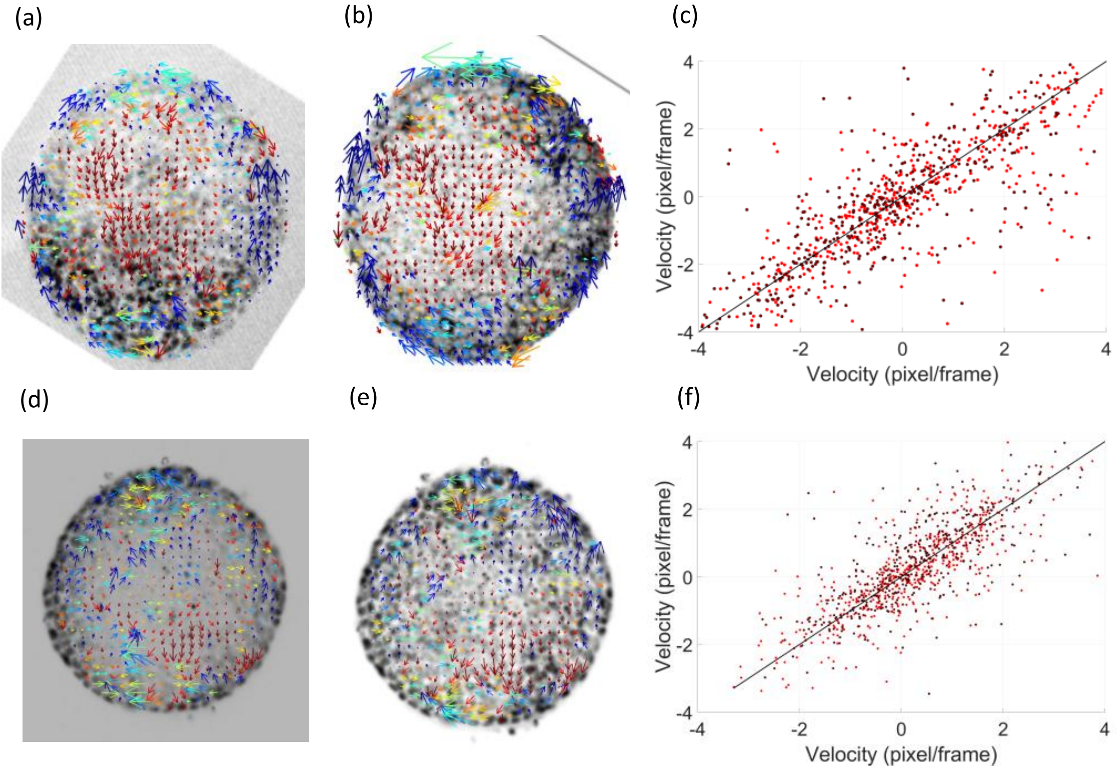


Figure S7: Variations in velocity fields depending on the nature of the signal used in the optical flow pipeline. (a-b) Raw velocity fields obtained between two consecutive time points separated by 5 minutes during symmetry breaking: (a) on the T/Bra GFP signal (with enhanced constrast to improve visibility of even low-T/Bra cells), and (b) on the Sulforhodamin B signal. (c) Comparison between the velocity components obtained in (a) and (b) with the x component of the velocity in red and the y component of the velocity in brown. The black line corresponds to equality between results from a and b. (d-e) Raw velocity fields obtained between two consecutive time points separated by 5 minutes at early symmetry breaking: (d) on the raw nuclei signal (SPY555-DNA), and (e) on the signal with a locally equalized contrast in order to improve visibility of the inner nuclei. (f) Comparison between the velocity components obtained in (d) and (e) with the x component of the velocity in red and the y component of the velocity in brown. The black line corresponds to equality between results from d and e.

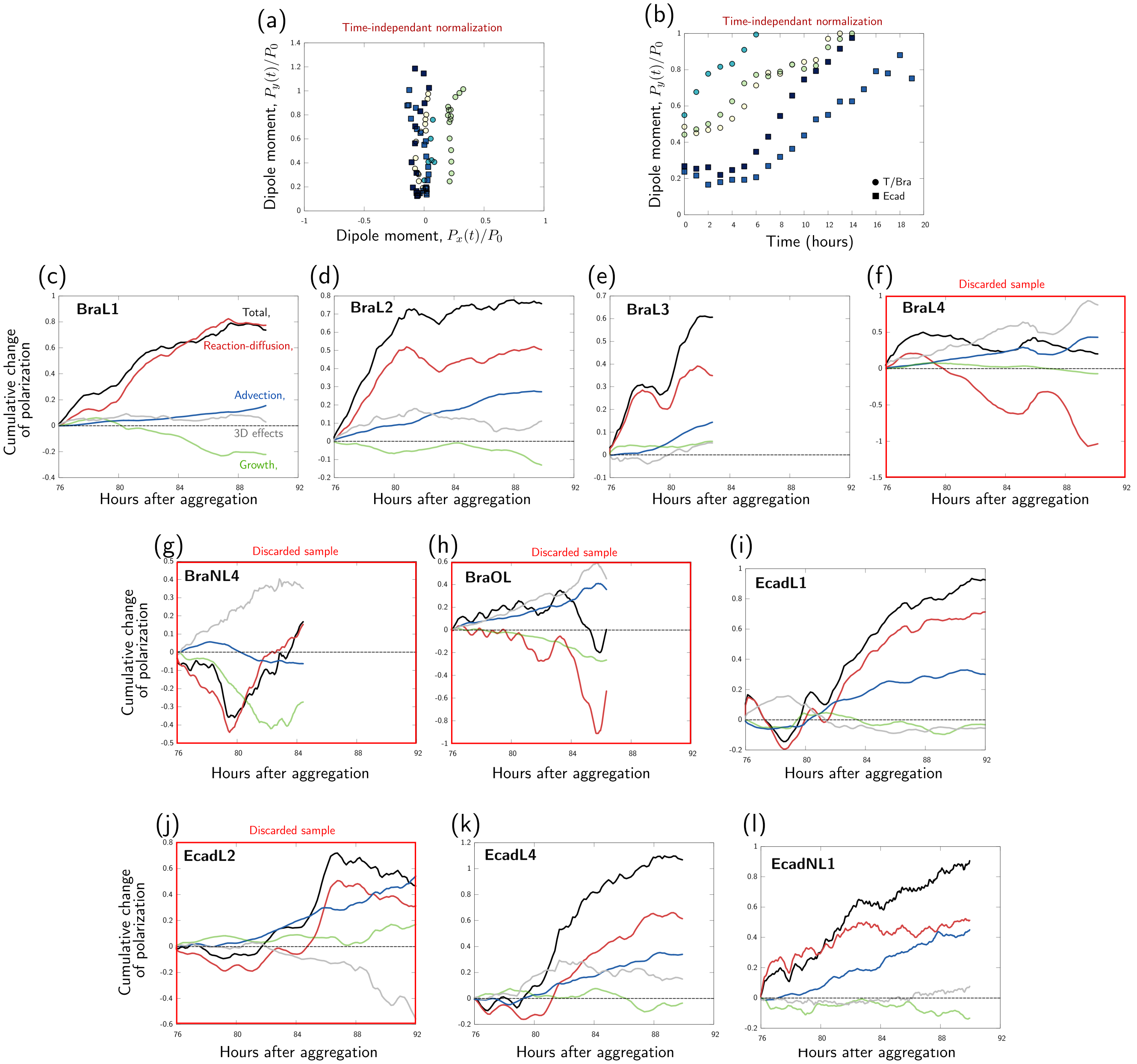


Figure S8: Dipole moment analysis. (a) Trajectories of the $x$ and $y$ components of the dipole moment $\boldsymbol{P} = (P_x, P_y)$, for different samples. The $y$ axis is defined as the main axis of elongation of the aggregate at the last time point. (b) $P_y$ component of the dipole moment over time, for different samples. (c-l) Cumulative contributions to the polarization for 10 gastruloids. These include the 6 samples considered in Figure 2 of the main text, while 4 gastruloids were discarded (red frames) because of a too large 3D contribution (see SI section Analysis procedure).

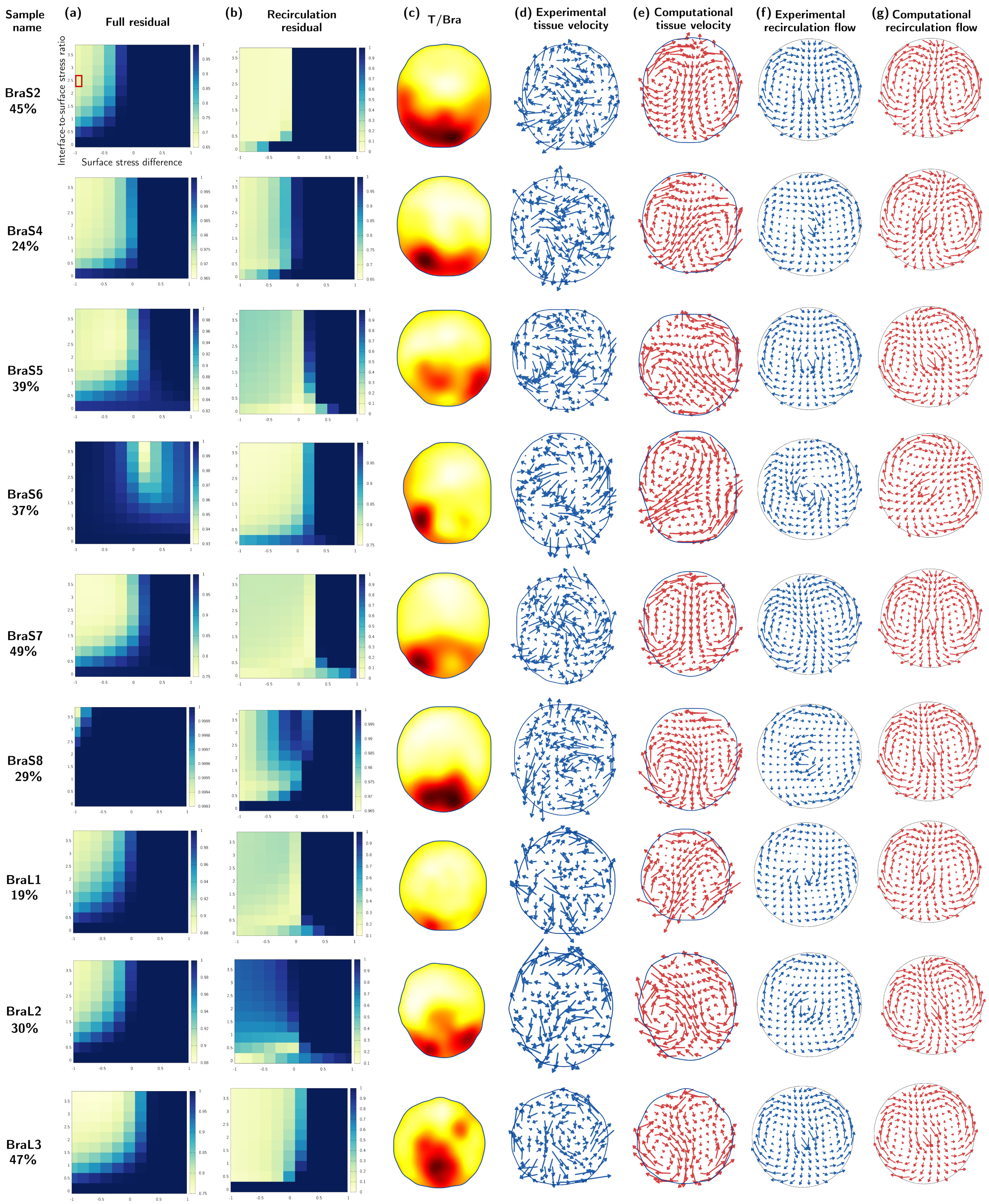


Figure S9: Simulations of tissue flows during symmetry breaking of 9 gastruloids. The first column indicates the sample name and the relative contribution of the recirculation mode to the experimental flow fields. For each sample, we show (a) the computational residual as a function of the surface stress difference $\Delta\kappa_s$ and the interface-to-surface stress ratio $r$, (b) the residual comparing the recirculation flow, Eq. (S23), of simulated and experimentally observed velocity fields (columns f and g), (c) the experimental T/Bra field, (d) the experimental tissue velocity field, (e) the computational velocity field for $r = 2.5$ and $\Delta\kappa_s = -1$, (f) the reconstructed experimental recirculation flow and (g) the reconstructed computational recirculation flow for $r = 2.5$ and $\Delta\kappa_s = -1$.

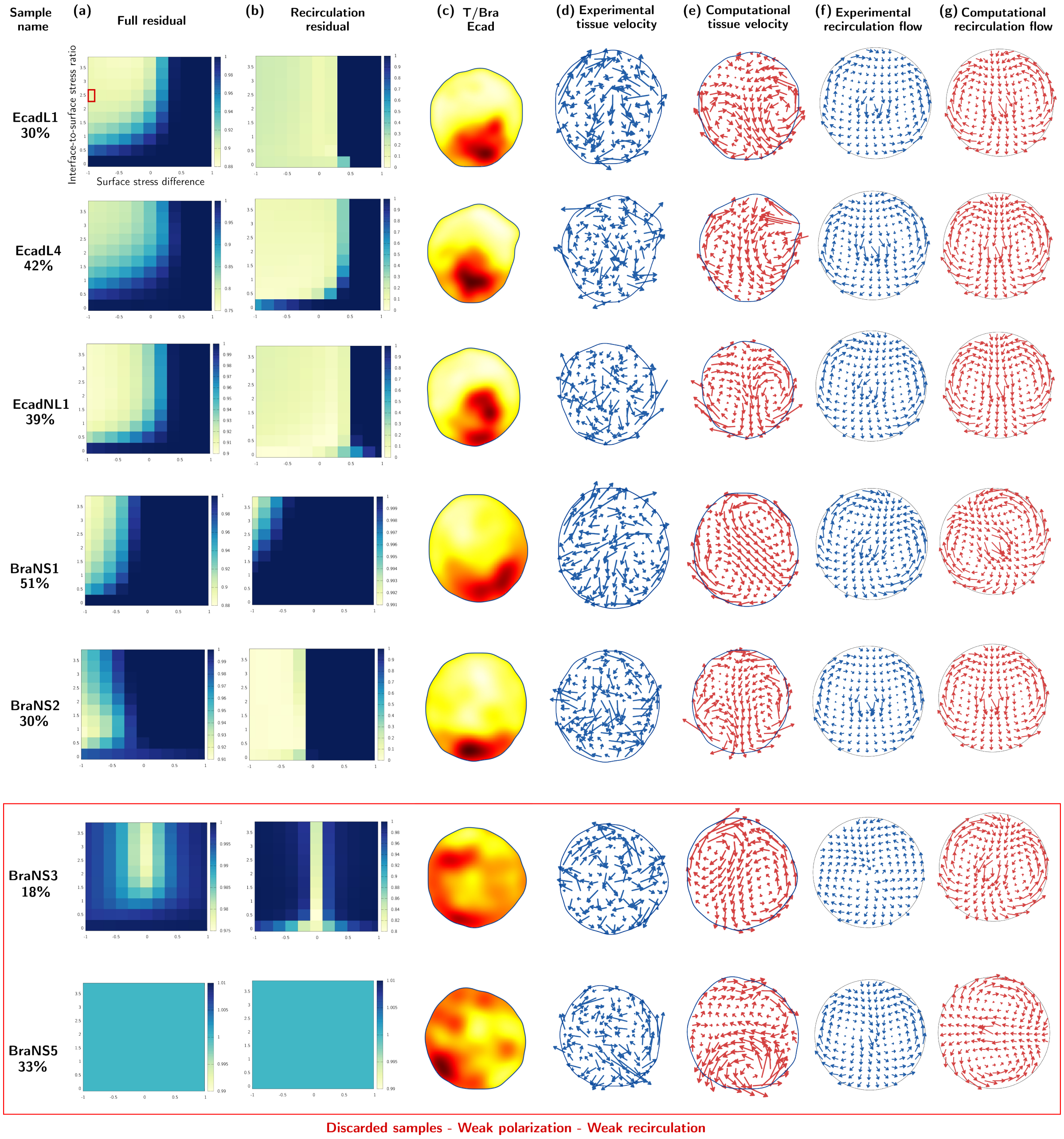


Figure S10: Simulations of tissue flows during symmetry breaking of 7 gastruloids. The first column indicates the sample name and the relative contribution of the recirculation mode to the experimental flow fields. For each sample, we show (a) the computational residual as a function of the surface stress difference $\Delta\kappa_s$ and the interface-to-surface stress ratio $r$, (b) the residual comparing the recirculation flow, Eq. (S23), of simulated and experimentally observed velocity fields (columns f and g), (c) the experimental T/Bra or Ecad field, (d) the experimental tissue velocity field, (e) the computational velocity field for $r = 2.5$ and $\Delta\kappa_s = -1$, (f) the reconstructed experimental recirculation flow and (g) the reconstructed computational recirculation flow for $r = 2.5$ and $\Delta\kappa_s = -1$.

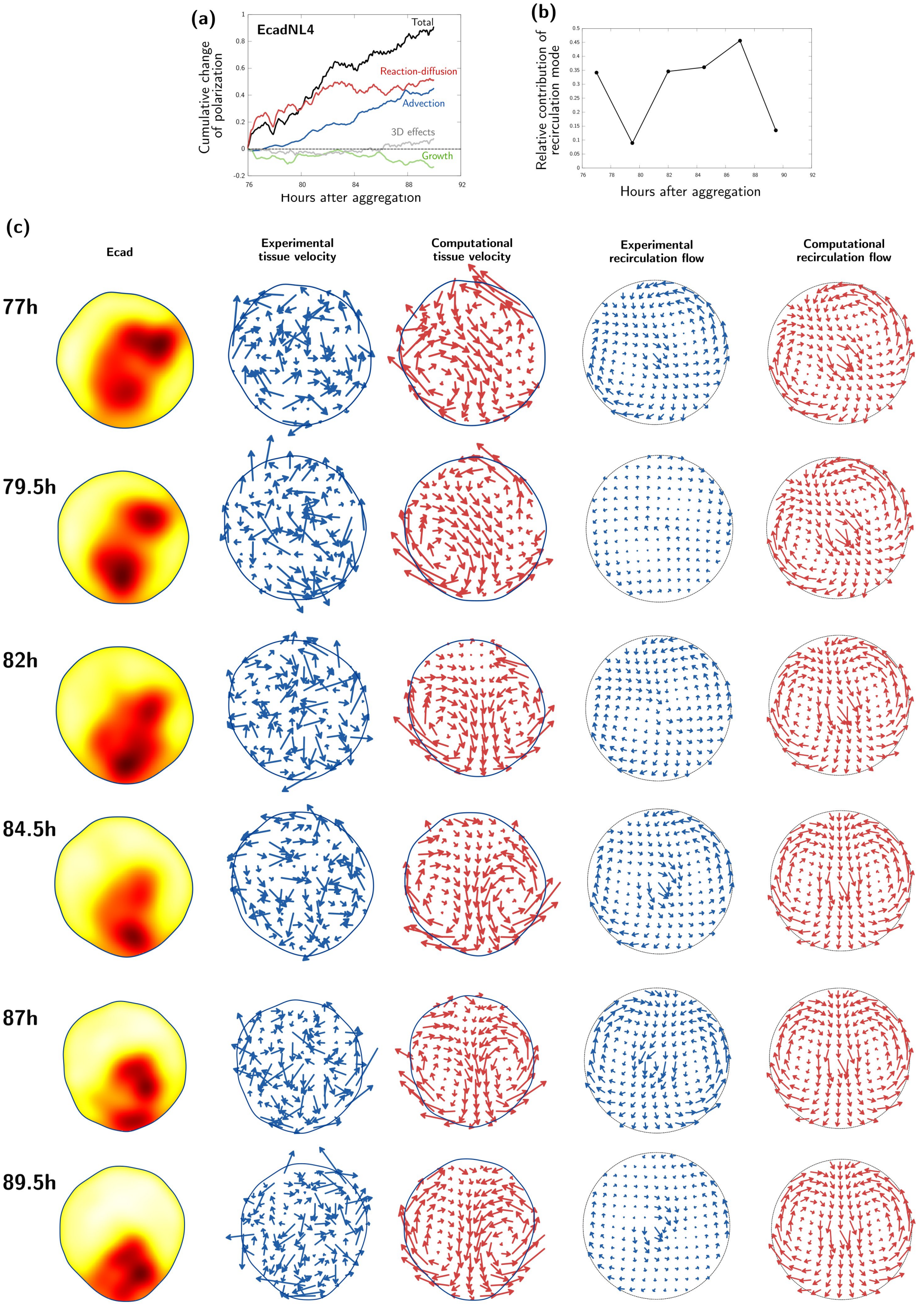


Figure S11: Tissue flows during symmetry breaking for one gastruloid with Ecad cell line, at different times. (a) Dipole moment decomposition. (b) Relative contribution of the recirculation flow in the experimental flow fields. (c) Ecad field, experimental and computational flow fields. Computational flows are computed for $r = 2.5$ and $\Delta\kappa_s = -1$.

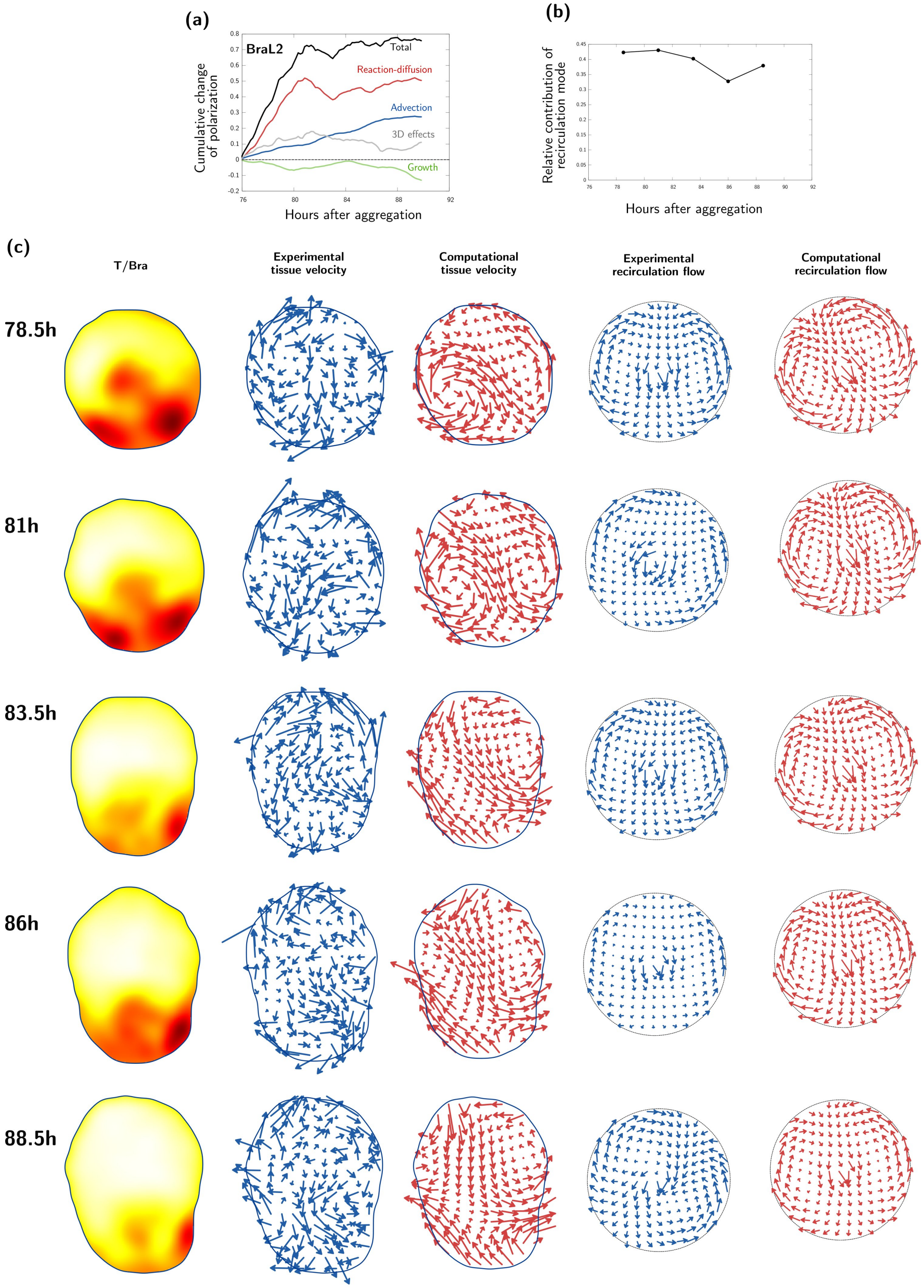


Figure S12: Tissue flows during symmetry breaking for one gastruloid with T/Bra cell line, at different times. (a) Dipole moment decomposition. (b) Relative contribution of the recirculation flow in the experimental flow fields. (c) T/Bra field, experimental and computational flow fields. Computational flows are computed for $r = 2.5$ and $\Delta\kappa_s = -1$.

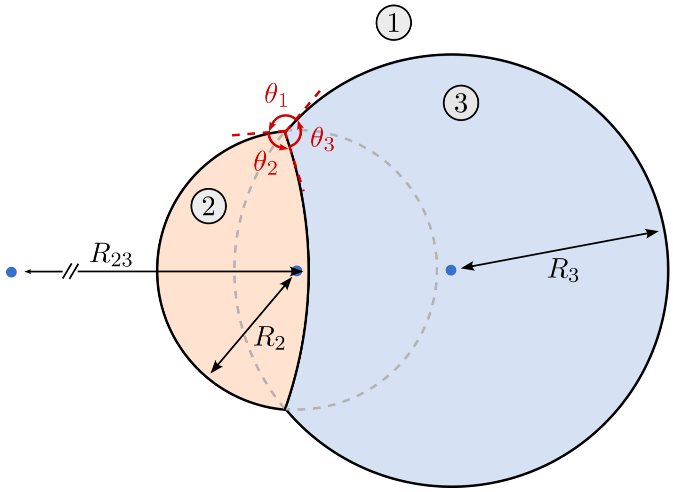


Figure S13: Schematic representation of the equilibrium between two immiscible droplets.

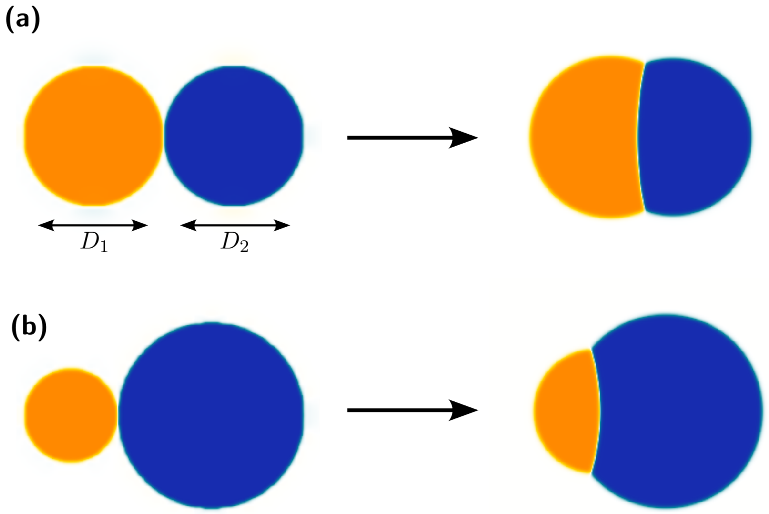


Figure S14: Simulation of fusing droplets with different diameter ratios $D_1/D_2$, for $r = 0.8$ and $\Delta\kappa_s = -0.2$. (a) $D_1/D_2 = 1$. (b) $D_1/D_2 = 1/2$.

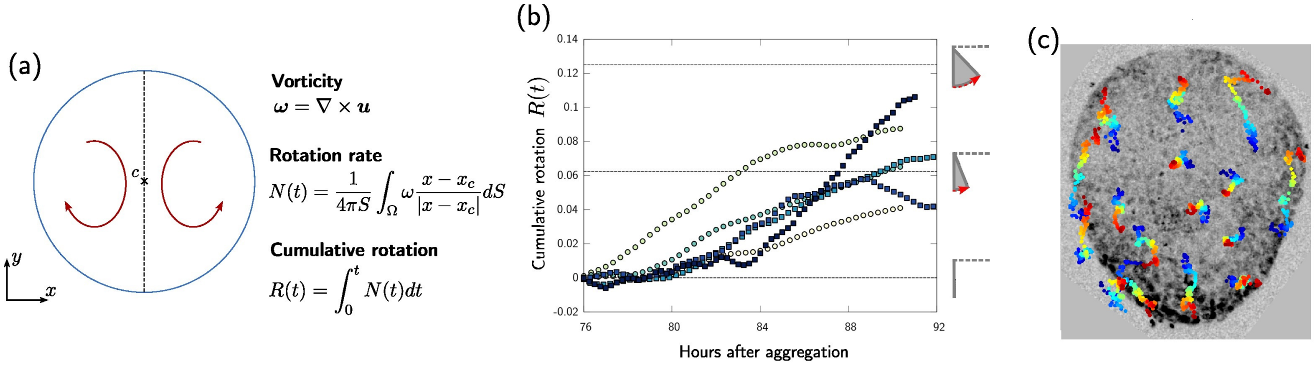


Figure S15: Quantification of the overall cell motion during symmetry breaking. (a) We use the coarse-grained tissue velocity $\boldsymbol{u}$ to determine the average tissue rotation rate $N(t)$ on both sides of the aggregate as well as the cumulative tissue rotation $R(t)$. (b) Over 6 samples, the average cumulative rotation is between 1/16 and 1/8 complete rotation by the end of the symmetry breaking process. The symbols and colors are the same as in Figure 2(c) in the main manuscript. (c) Even when the computed average rotation is small, in some regions cells can move over approximately half of the aggregate length during polarization, as shown by selected trajectories computed using the coarse-grained tissue flow. This sample corresponds to the white circle symbols in (b). On trajectories, blue and red indicate initial and final position, respectively.

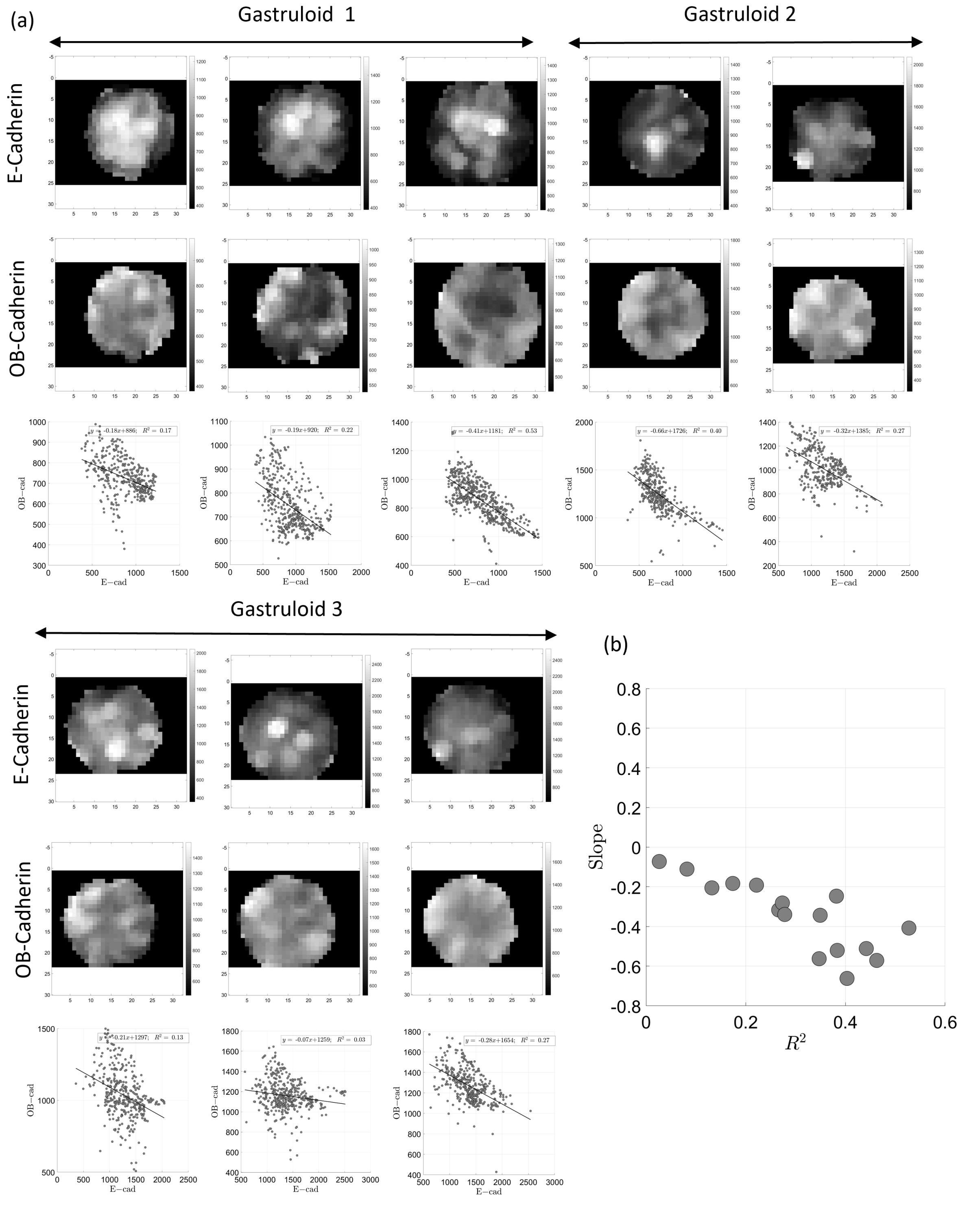


Figure S16: Spatial anti-correlation of E-cadherin and OB-cadherin (Cadherin 11). (a) Immunostaining of E-cadherin and OB-cadherin were performed for 3 different 76hAAF gastruloids (see Methods, Immunostainings, and Supp Movie OB-Ecad). Each image corresponds to a coarse-grained intensity (CGI) map of cadherins (each pixel in the map is spaced by 7 µm and its intensity corresponds to the raw image intensity averaged in a disk of radius 14 µm). Each raw image is obtained by averaging 10 µm-thick stacks (10 slices) at different z locations in the gastruloid. CGI values of OB-cadherin are ploted as a function of CGI values of E-cadherin in the same place and fitted linearly. (b) Slopes of the linear fits in panel (a) as a function of their respective $R^2$ values.

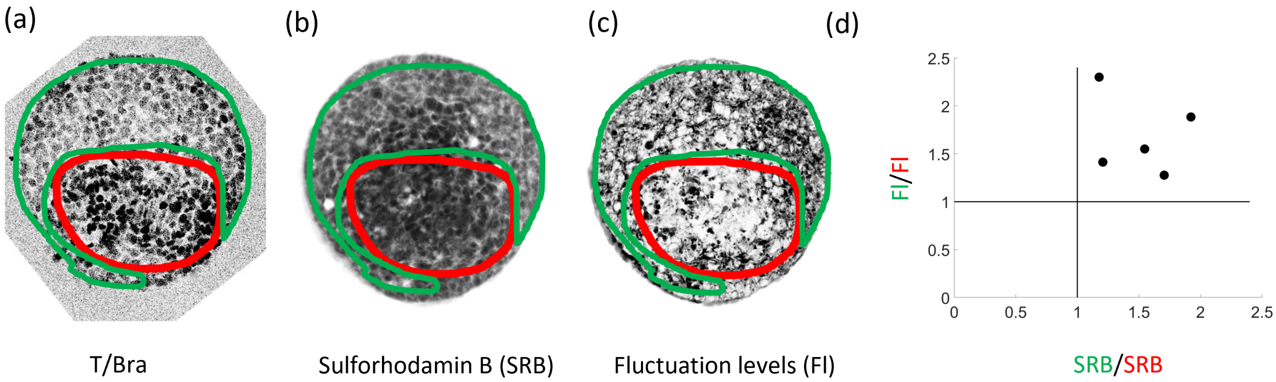


Figure S17: At a coarse-grained level, T/Bra expression is positively correlated with tissue packing density and negatively correlated with cell-scale motion. (a) For a gastruloid mid-plane at 80hAAF, the T/Bra signal (inverted) is used to manually divide the cross-section into a T/Bra positive region (red) and a T/Bra negative region (green). (b,c) Regions as defined in panel a superimposed on the Sulforhodamin B signal, which indicates inter-cellular space. (b) Average ("SRB") over one hour (12 frames), which we use as a proxy for cell packing density. (c) Variance ("Fl") over one hour (12 frames), which we use as a proxy for cell-scale motion. (d) Ratios of both SRB and Fl as defined in panels (b) and (c) between the T/Bra negative region (green) and the T/Bra positive region (red). Both SRB and Fl are higher in the T/Bra negative region indicating a lower cell packing density and a higher cell motility.

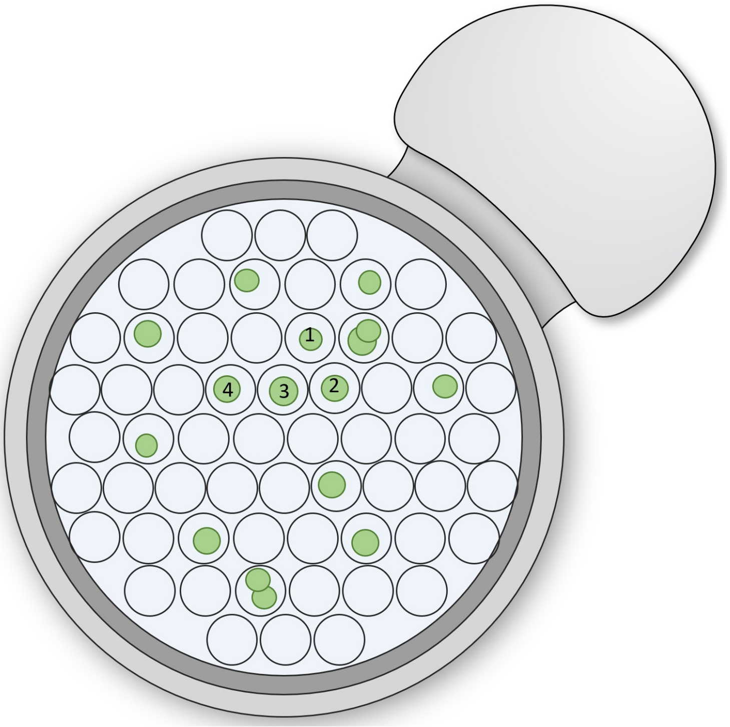


Figure S18: Gastruloids random seeding in the microwells system (SUNBIOSCIENCE ref: *Gri3D*). 72 hAAF gastruloids are seeded in the system. Each well has zero, one or multiple gastruloids after the random seeding. A serie of 3 or 4 neighbouring gastruloids is chosen (as the ones numbered from 1 to 4 in the schema) for parallel imaging, in order to minimize the distance covered with the water objective to avoid bubbles emergence over time.
